# Confined T cell migration controls programmed cell death 1 expression

**DOI:** 10.64898/2026.07.03.736145

**Authors:** Yunus Alapan, Jaehoon Kim, Kiyoon Min, Alexander Shih, Samuel N. Lucas, Taehee Yoon, Paul A. Archer, Heather K. Lin, Ruby Freeman, Megen C. Wittling, Megan M. Wyatt, Chrystal M. Paulos, Sarwish Rafiq, Susan N. Thomas

## Abstract

T cell immune checkpoint expression and dysfunction are attributed solely to molecular cues. We discover using microphysiological systems and *in vivo* models that transmigration through confined pores induces acute loss of programmed cell death 1 within minutes of surface immune checkpoint markers on CD8+ T cells through ubiquitin-mediated proteasomal degradation, corresponding with enhanced fitness and function. Such imprints and its underlying mechanism of programmed cell death 1 loss are conserved across species and applicable to human TIL and CAR T cells, and are correlative with disease outcomes in human melanoma. Our findings establish the diverse tissues landscapes that T cells traverse during immunosurveillance, and specifically transmigration, as a form of mechanical immune regulation, and reveal a mechano-modulatory strategy to improve the quality and persistence of engineered or patient-derived T cells for adoptive immunotherapy.

## Introduction

Immune surveillance of the human body depends on the capability of T cells to traverse widely varying biophysical microenvironments across numerous tissues and organs (*1*). As T cells circulate, they encounter not only biochemical cues such as antigens and cytokines but also a range of physical barriers—including vascular endothelium, basement membranes, and dense extracellular matrix—that must be overcome to enter target tissues. Protein expression and biophysical profiles required for T cells to engage successive microenvironments are shaped by remodeling events induced by prior molecular and cellular interactions.

Recent studies suggest that biophysical cues within tissues can modulate T cell function through mechanosensitive pathways. At the cellular level, T cell-exerted forces are required for cytotoxic killing of target cells (*2*) and cancer cells exploit this phenomenon to evade killing by decreasing their membrane stiffness (*2–4*). Higher stiffness of extracellular matrix in solid tumors has been linked to increased expression of programmed cell death (PD) 1 and Tim3, suggesting exhaustion, induced through mechanosensory pathways within T cells (*5*). PD-1 is widely perceived as a stable imprint of antigenic history. Yet, rapid post-translational, via recycling, shedding, and proteasomal degradation (*6–8*) reveals that its expression is far more dynamic, shaped by non-transcriptional cues.

These findings suggest that biophysical interactions with distinct microenvironments can dynamically tune T cell function, acting as either inhibitory or stimulatory checkpoints. Trans-endothelial migration, a ubiquitous step for cellular entry into solid tissues, forces active movement through highly confined spaces under adhesive and chemokine guidance. Yet whether this mechanical process rewires functional states or surface checkpoint expression has remained unexplored. Here, we uncover how transmigration mechanistically reprograms T cell immune checkpoint expression, revealing a previously unrecognized layer of immune regulation.

## Results

### Transmigration through confined channels in vitro and in vivo triggers acute PD-1 loss in CD8^+^ T cells

We engineered a microfluidic transmigration on a chip (T-Chip) platform that recapitulates the key physiological processes of T cell homing to tissues from the circulation (Fig. 1A and fig. S1, A and B): selectin-mediated rolling, integrin-based firm adhesion, and chemokine-guided transmigration (*1*). The microporous membrane (3 µm pore diameter) mimicking endothelial gaps generated during T cell transmigration (*9*) is functionalized with adhesion molecules (E-and P-selectin, ICAM-1, VCAM-1) and positioned to physically separate the upper and lower channels of the T-Chip, enabling selective recovery of transmigrated cells (Fig. 1B and fig. S1B). The branched upper channel ensures uniform distribution of physiologic wall shear stress (0.75 dyn/cm^2^) (*10*) over the membrane surface (fig. S1C). The lower channel contains a chemokine cocktail (CCL2-5 and CXCL9/10) that mimics the transmigration axis of inflamed tissues and tumors (*11–13*). Uniform chemokine gradients are generated across the membrane (fig. S1D) with defined pore densities (6×10^5^ or 2×10^6^ pores/cm²), facilitating transmigration within minutes by murine CD8^+^ T cells expanded ex vivo with Dynabeads for 10 days (Fig. 1C, and movie S1).

**Figure 1.**
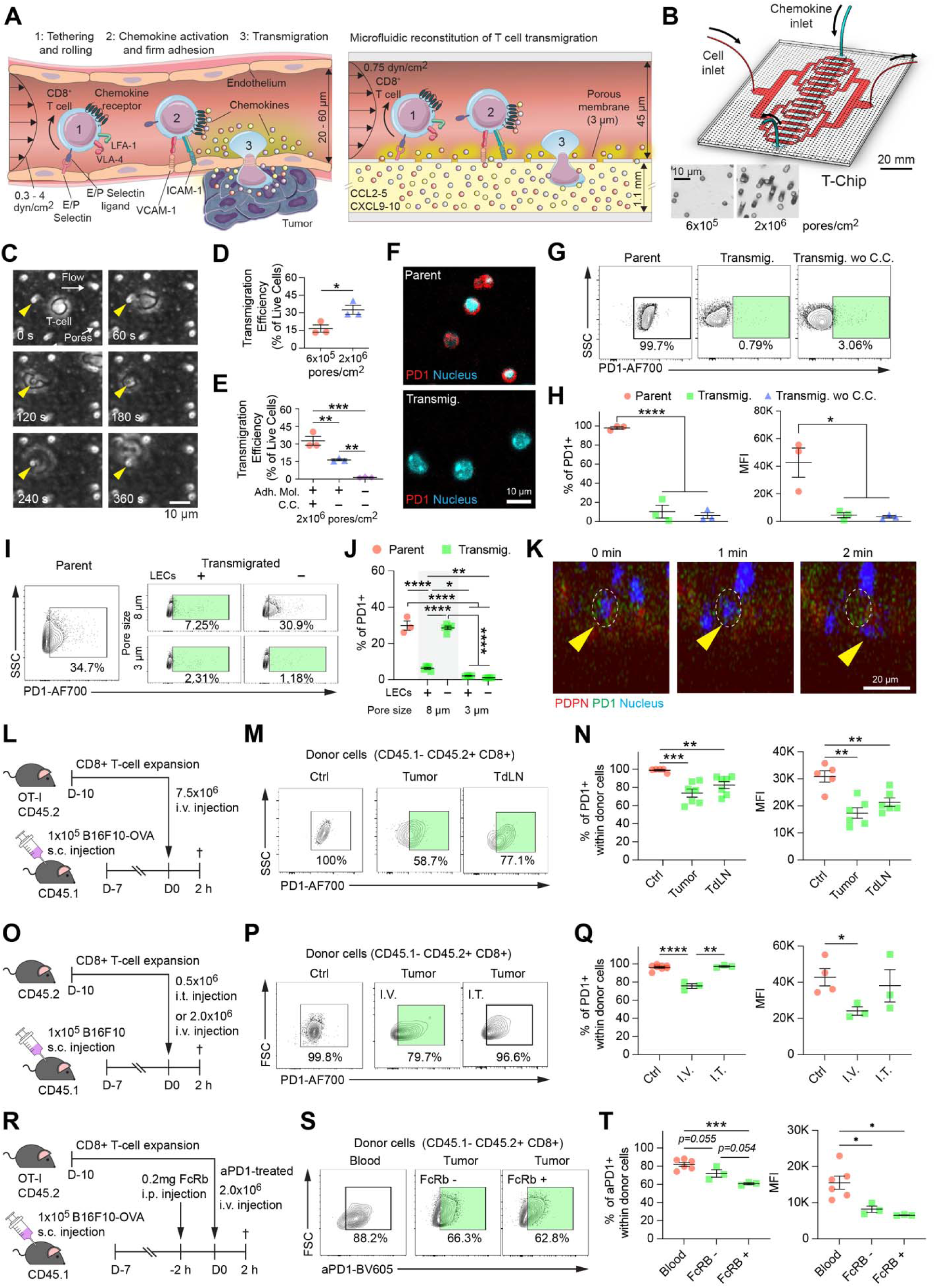
Transmigration induces acute loss of surface PD-1 on CD8+ T cells in vivo and in vitro. (A) Microfluidic reconstitution of T cell transmigration in vitro. Physiological steps of T cell transmigration in the tumor microenvironment (left): (1) rolling and tethering via adhesion molecules, (2) firm adhesion to the endothelium, and (3) chemokine-induced transmigration into the tumor. Design of microfluidic transmigration-on-a-chip (T-Chip), recapitulating physiological processes in vitro (right). Expanded mouse CD8+ T cells are perfused over a porous membrane under physiological shear stress (0.75 dyn/cm2), and allowed to adhere on surface-coated adhesion molecules (E/P selectin, ICAM-1, VCAM-1) and transmigrate through micron sized pores under the guidance of chemokines (CCL2-5, and CXCL9/10) supplied in the lower channel. (B) 3D schematic of T-Chip (top) incorporating parallel upper channels for cell perfusion and a lower channel as a chemokine reservoir. The upper and lower channels are separated by a porous membrane with 3 µm diameter pores (scale bar: 20 mm). Bright field images of microporous membranes (bottom) with low (6×10^5^ pores/cm^2^) and high (2×10^6^ pores/cm^2^) pore densities (scale bar: 10 µm). (C) Representative time-lapse bright-field images of an individual mouse T cell transmigrating through a pore. Yellow arrowheads indicate the pore location. (scale bar: 10 µm). (D) Membranes with greater pore density (2×10^6^ vs 6×10^5^ pores/cm^2^) enabled enhanced transmigration efficiency (% of perfused live cells collected from the lower channel) (n = 3 chips per group; mean ± SEM; *p < 0.05, unpaired t-test). (E) Reduction in transmigration efficiency in the absence of only surface-coated adhesion molecules (Adh. Mol.) or both adhesion molecules and chemokine cues (C.C.) (n = 3 chips per group; mean ± SEM; **p < 0.01, ***p < 0.001, one-way ANOVA with Tukey’s multiple comparison test). (F) Immunofluorescence confocal microscopy imaging revealed substantial decrease of PD-1 expression (red) on transmigrated CD8+ T cells recovered from the T-Chip compared to their non-perfused parent population. Cyan shows nucleus. (scale bar: 10 µm) (G) Representative flow cytometry plots for PD-1 expression on parent cells, transmigrated cells, and transmigrated cells without chemokine cues (Transmig. wo C.C.). Green gates highlight the shift toward reduced PD-1 expression in transmigrated cells relative to the parent population. Full gating strategy for flow cytometry may be found in fig. S2A. (H) % PD-1+ cells (left) and PD-1 median fluorescence intensity (MFI, right) were significantly lower for both transmigrated cells with and without chemokine cues, indicating chemokine-independent downregulation of PD-1. (n = 3 chips per group; mean ± SEM; ****p < 0.0001, *p < 0.05, one-way ANOVA with Tukey’s multiple comparison test) (I) Representative flow cytometry plots of PD-1 expression in parent cells and transmigrated cells recovered from transwells with or without an LEC monolayer using 8 µm or 3 µm pore membranes. Green gates highlight the PD-1-low population observed in transmigrated cells. (J) Quantification of PD-1+ frequencies across the indicated conditions. Across 8 µm pores, PD-1 downregulation was observed only when transmigration occurred in the presence of an LEC monolayer, whereas membrane-only controls did not elicit PD-1 loss. Across 3 µm pores, PD-1 downregulation was observed in transmigrated cells regardless of LEC presence. (n = 3-5 wells per group; mean ± SEM; one-way ANOVA with Tukey’s multiple comparisons test). (K) Live confocal imaging of T-cell transmigration across an LEC monolayer in a Transwell insert. LECs were labeled with podoplanin (PDPN), and CD8⁺ T cells were pre-labeled for PD-1. The time-lapse images show show a PD-1–labeled T cell (arrowhead; dashed outline) undergoing transmigration through an LEC monolayer, accompanied by a progressive reduction in PD-1 signal during transit. (K) Experimental design of in vivo adoptive transfer experiment for short-term T cell homing. CD8+ T cells were isolated from CD45.2+ OT-I mice, expanded ex vivo for 11 days, and intravenously (i.v.) injected into B16F10-OVA tumor-bearing CD45.1+ mice (1×10^5^ tumor cells implanted subcutaneously 7 days prior). Tumors and tumor-draining lymph nodes (TdLNs) were harvested at 2 h post-injection for analysis. (L) Representative flow cytometry plots for PD-1 expression on donor CD8+ T cells (CD45.1-CD45.2+) isolated from tumors and TdLNs at 2 h post-transfer, compared to non-injected cells, which indicate control group (Ctrl). Green gates highlight donor T-cell populations with reduced PD-1 expression in tumors and TdLNs relative to the control population. Full gating strategy for flow cytometry may be found in fig. S5B. (M) Summary of PD-1 frequency and median fluorescence intensity (MFI) on donor CD8+ T cells recovered from tumors and TdLNs 2 h after transfer. Donor T cells recovered from both tissues exhibited reduced PD-1 expression compared with the pre-transfer control (Ctrl) population. (n = 5-7 mice per group; mean ± SEM; **p < 0.01, unpaired t-test) (N) Experimental schematic comparing intratumoral (i.t.) versus intravenous (i.v.) delivery of expanded donor CD8+ T cells into B16F10 tumor-bearing mice, followed by analysis 2 h after transfer. (O) Representative flow cytometry plots showing PD-1 expression on donor CD8⁺ T cells recovered from tumors following i.t. or i.v. delivery. Green gates highlight donor T-cell populations with reduced PD-1 expression relative to the pre-transfer control population. (P) Quantification of PD-1 frequency and median fluorescence intensity (MFI) of donor CD8+ T cells recovered from tumors after i.t. versus i.v. delivery. PD-1 downregulation was observed following i.v. transfer, whereas i.t. delivery did not induce PD-1 loss at the same time point. (n = 3-8 mice per group; mean ± SEM; ****p < 0.0001, **p < 0.01, *p < 0.05, unpaired t-test) (Q) Experimental schematic of in vivo PD-1 tracking. Expanded OT-I CD45.2+ CD8+ T cells were pre-labeled with a fluorescently conjugated anti–PD-1 antibody and intravenously (i.v.) transferred into CD45.1+ B16F10-OVA tumor-bearing mice. Recipient mice received Fcγ receptor (FcγR) blocking reagent by intraperitoneal (i.p.) injection 2 h before transfer to minimize FcγR-mediated antibody stripping. Tumors were harvested 2 h after transfer for analysis of donor T cells. (R) Representative flow cytometry plots and **(S)** quantification of anti–PD-1 signal frequency and median fluorescence intensity (MFI) of donor CD8⁺ T cells (CD45.1- CD45.2+ CD8+) recovered from blood and tumor at 2 h post-transfer. Green gates highlight donor T-cell populations with reduced pre-bound anti–PD-1 signal in tumors relative to blood. The frequency and MFI of anti–PD-1+ donor cells were reduced in tumor relative to blood, indicating loss of pre-bound anti–PD-1 signal after tissue entry. (n = 3-6 mice per group; mean ± SEM; ***p < 0.001, *p < 0.05, unpaired t-test).

When expanded CD8^+^ T cells were perfused over membranes with 2×10^6^ pores/cm^2^, >30% of cells transmigrated within 3 h, whereas membranes with 6×10^5^ pores/cm^2^ yielded ∼15% efficiency (Fig. 1D). Removing chemokines from the lower channel reduced transmigration by nearly half. Further removal of adhesion molecules completely abrogated transmigration, indicating an essential role for adhesion in this process (Fig. 1E) (*14*). Selective enrichment of live expanded CD8^+^ T cells was achieved by transmigration in T-Chip (fig. S2B) that maintained their capacity to expand upon ex vivo restimulation (fig. S2C), indicating that the process may inherently filter for viable or functional cells with high expansion capabilities and exclude apoptotic or damaged populations. Memory subset profiling of perfused T cells based on CD44 and CD62L expression via multicolor flow cytometry revealed increased effector memory (TEM) and decreased naïve fractions in transmigrated cells (fig. S3, A and B). However, similar phenotypic changes have been observed in T cells interacting with selectin-coated microfluidic channel surfaces, suggesting that adhesion molecule signaling may contribute to CD44 and CD62L modulation (*14*). T-Chip thus facilitates efficient T cell transmigration with high fidelity to yield large quantities of viable, effector memory polarized cells.

Transmigrated CD8^+^ T cells recovered from T-Chip comprised of either 3 or 5 µm pores exhibited a profound reduction in the frequency of PD-1^+^ cells and PD-1 median fluorescence intensity compared to their non-perfused parent T cell counterpart, as demonstrated by confocal microscopy and confirmed by flow cytometry (Fig. 1F-H fig. S2D). Notably, a similar degree of PD-1 loss was observed even in the absence of chemokines, suggesting that the physical act of transmigration, rather than chemokine-mediated signaling, is the key mediator of this effect, and occurred across all memory phenotypes, suggesting that its loss is not simply a consequence of differentiation status (fig. S2E). Transmigrated CD8^+^ cells also exhibited reduced CXCR3 and CD62L, along with elevated CD44 and Ki-67 in comparison to non-perfused parent cells, parent cells exposed to the same chemokine cocktail in static conditions, and non-migrated cells that were perfused but did not transmigrate (fig. S3, C and D). In contrast, Granzyme B expression was similar across groups. Notably, although both PD-1 and CXCR3 were downregulated in transmigrated CD8^+^ T cells, their regulatory mechanisms may differ. In contrast to PD-1, which was downregulated regardless of chemokine absence (Fig. 1H), CXCR3 was less downregulated when CD8^+^ T cells transmigrated in the absence of chemokines (fig. S4A), suggesting that CXCR3 downregulation is more dependent on ligand engagement. Supporting this, CXCR3 was also modestly reduced in T cells exposed to chemokines under static conditions (fig. S3D), indicating that chemokine signaling alone induces partial receptor internalization (*15*). CXCR3 also rapidly recovered within 24 h post-transmigration (fig. S4B), indicating a transient response. This downregulation may reflect desensitization to chemokine signaling for transient tissue retention.

To evaluate whether transmigration induced changes in PD-1 expression resulting from T-Chip processing are recapitulated in conventional, field-standard assays, PD-1 expression was evaluated in in vitro Boyden chamber assays using Transwell inserts. Using a similar chemokine cocktail, PD-1 levels were observed to decrease in CD8^+^ cells transmigrated through Transwell inserts with 3 but not 8 µm pores, reflecting results obtained from T-Chip (Fig. 1, I and J) and suggesting an effect of confinement on this biological response to transmigration. However, the effect of PD-1 down regulation was recovered for 8 µm in the presence of lymphatic endothelial cell (LEC) monolayers, suggesting physiological barriers to transmigration can induce this biologic response. A progressive loss of PD-1 signal during CD8+ T cell transit was also revealed by in situ visualization of PD-1–labeled CD8+ T cells undergoing transmigration across an LEC monolayer in a Transwell insert enabled by live confocal imaging also revealed a progressive loss of PD-1 signal during transit (Fig. 1K).

To address whether PD-1 loss occurs in vivo during early tumor infiltration, PD-1 expression by OT-I CD8^+^ T cells activated via in vitro expansion over 10 d prior to transfer recovered early 2 h post transfer, and thus before the onset of chronic antigen exposure and immunosuppressive signaling, via intravenous injection into B16F10 tumor-bearing mice (CD45.1^+^) was interrogated (Fig. 1L). Whereas PD-1 on donor CD8^+^ T cells (CD45.2^+^) from blood showed only a modest reduction compared with that of the pre-transferred population (fig. S5C), donor T cells recovered from tumors and tumor-draining lymph nodes (TdLNs) exhibited a marked reduction in surface PD-1 expression (Fig. 1, M and N), as compared to those recovered at 24 h (fig. S5D), an effect not seen in the spleen (fig. S5) and that occurred irrespective of tumor-expression of cognate antigen (fig. S6). Consistent with PD-1 loss being triggered by tumor entry-associated, acute PD-1 downregulation occurred only after extravasation from the circulation after donor cell infusion into the blood, as transfer via an intratumoral administration resulted in no loss of PD-1 signal (Fig.1 O-Q). Notably, OT-I CD8^+^ T cells activated via in vitro expansion over 10 d prior prelabeled with fluorescent anti-PD-1 mAb recovered from tumors 2 h after transfer exhibited a loss of pre-bound anti–PD-1 (Fig.1 R-T). Tissue infiltration by T cells from the circulation thus downregulates PD-1 expression to influence the extent of therapeutic immune checkpoint inhibition.

Together, these results establish the T-Chip as a tractable system for scalable recovery of transmigrated cells at yields consistent with application to a variety of downstream applications (fig. S7) that recapitulates PD-1 down-regulation observed conventional physiological transmigration in vitro models as well as in vivo in syngeneic mouse tumor models by CD8^+^ T cells during early tumor infiltration.

### Transmigrated CD8^+^ T cells exhibit augmented multi-potency, cytotoxicity, proliferation, migration, and *in vivo* tumor control

Effects of transmigration-induced phenotypic remodeling of murine CD8^+^ T cells on cytotoxicity, proliferation, morphology, migration, and *in vivo* tumor control were assessed. Transmigrated CD8^+^ T cells produced significantly higher IFN-γ and TNF-α compared to parent cells following antigenic stimulation under co-culture with B16F10-OVA tumor cells, regardless of tumor PD-L1 status induced by IFN-γ pretreatment (Fig. 2, A and B), whereas IL-2 production was only modestly increased. Transmigrated OT-I CD8^+^ T cells also killed B16F10-OVA tumor cells more effectively than parent cells, regardless of IFN-γ pre-treatment or anti-PD-1 treatment (Fig. 2C). Notably, IFN-γ pre-treatment, which elevated PD-L1 expression on tumor cells (fig. S8A), impaired the cytotoxicity of parent cells but had minimal effect on transmigrated cells. However, anti-PD-1 treatment attenuated the cytotoxicity gap, suggesting that the enhanced cytotoxicity of transmigrated cells is mediated, at least in part, by their reduced PD-1 levels. During co-culture with tumor cells, proliferation potential was also augmented in transmigrated cells (Fig. 2D). Over 70% of transmigrated cells entered division, compared to fewer than 50% for parent samples, irrespective of the treatment condition. Transmigration thus primes CD8^+^ T cells for enhanced effector function, as evidenced by increased cytokine production (*16*), as well as elevated activation markers, memory polarization, proliferation and acute PD-1 loss.

**Figure 2.**
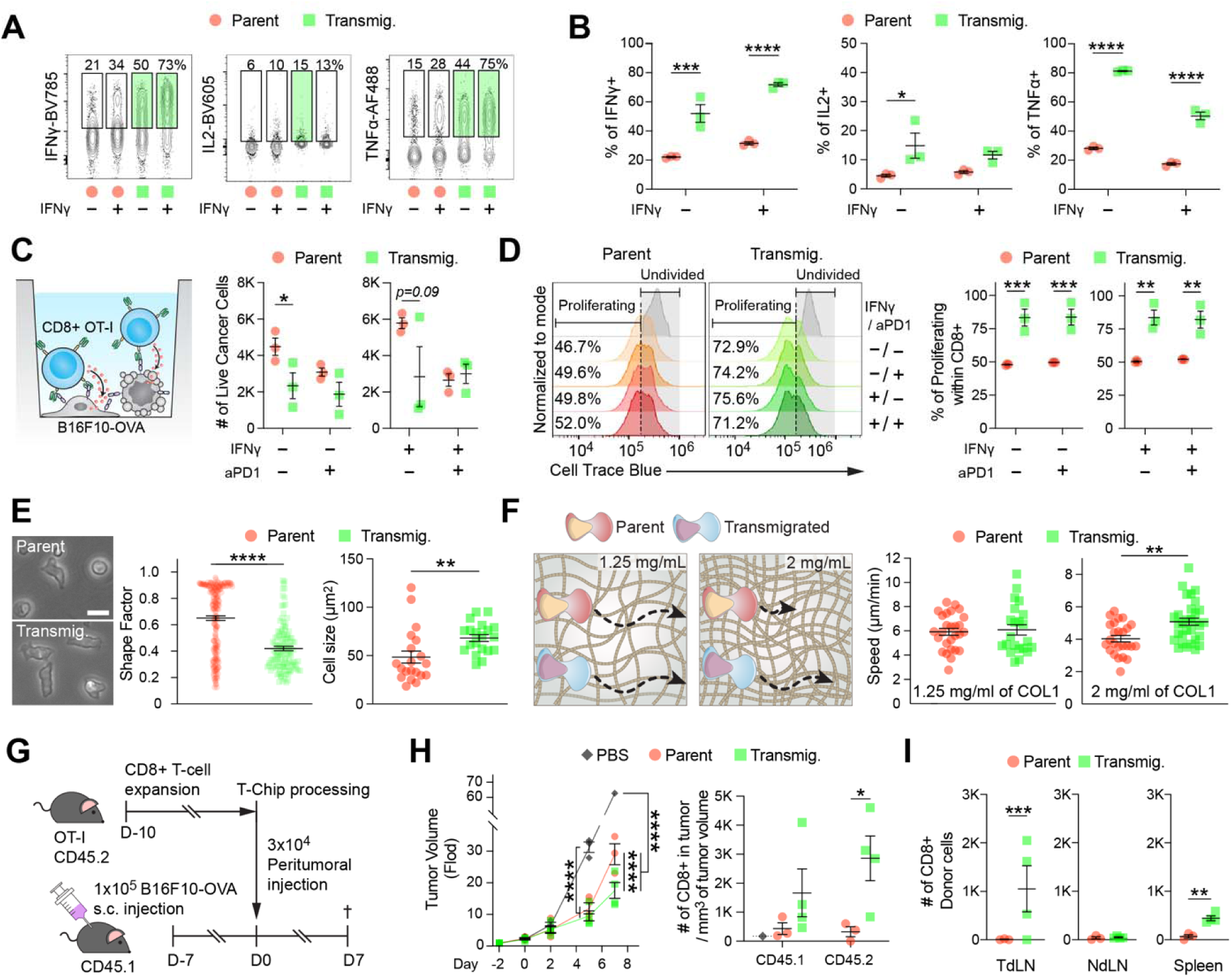
Transmigrated CD8^+^ T cells exhibit augmented multi-potency, cytotoxicity, proliferation, migration, and *in vivo* tumor control. **(A)** Representative flow cytometry plots for intracellular cytokine staining (IFN-γ, IL-2, and TNF-α) in parent and transmigrated OT-I CD8+ T cells after 3 h co-culture with B16F10-OVA cells, pre-treated with or without IFN-γ (10 ng/ml), in the presence of Brefeldin A (5 µg/mL). Green gates highlight cytokine-positive populations enriched following transmigration. Full gating strategy for flow cytometry may be found in fig. S8B. **(B)** Summary of cytokine-producing cells from (A). Transmigration significantly enhanced IFN-γ, IL-2, and TNF-α production. (n = 3–4 wells per group; mean ± SEM; *p < 0.05, ***p < 0.001, ****p < 0.0001, one-way ANOVA with Tukey’s multiple comparison test). **(C)** Schematic illustration (left) and quantification (right) of tumor cell killing by parent or transmigrated OT-I CD8⁺ T cells after 24 h co-culture with B16F10-OVA cells treated with or without IFN-γ (10 ng/ml) or anti PD-1 (aPD1, 10 µg/ml). (n = 3 wells per group; mean ± SEM; *p < 0.05, one-way ANOVA with Tukey’s multiple comparison test). **(D)** Proliferation profiles (left) and quantification (right) of parent and transmigrated OT-I CD8+ T cells using CellTrace Blue labelling after 24 h co-culture with B16F10-OVA tumor cells in the presence or absence of IFN-γ (10 ng/ml) and/or aPD-1 (10 µg/ml). Transmigrated cells showed increased proliferation regardless of IFN-γ or PD-1 blockade. (n= 3 wells per group; mean ± SEM; **p < 0.01, ***p < 0.001, one-way ANOVA with Tukey’s multiple comparison test) Full gating strategy for flow cytometry may be found in fig. S8B. **(E)** Representative bright-field images of parent and transmigrated T cells (left). Quantification (right) of shape factor and cell size, indicating polarization and enlargement of T cells after transmigration. (n > 100 cells per group, more than 3 experiments; mean ± SEM; ****p < 0.0001, **p < 0.01, unpaired t-test). **(F)** Schematic diagram of migration assay (left) and quantification of cell migration speed (right) in 3D collagen gels (1.25 and 2 mg/mL). Transmigrated cells migrated faster, particularly in 2 mg/ml collagen gel (n > 20 cells per group, more than 4 experiments; mean ± SEM; **p < 0.01, unpaired t-test). **(G)** Experimental design of in vivo peritumoral adoptive transfer experiment. OT-I CD8+ T cells expanded ex vivo for 11 days were processed in T-Chip and injected peritumorally into B16F10-OVA-bearing mice (1×10^5^ tumor cells implanted subcutaneously 7 days prior). Tumors were harvested for analysis 7 days post-injection. Full gating strategy for flow cytometry may be found in fig. S5B. **(H)** Tumor volume over time (Fold change from initial volume, left) and quantification of tumor-infiltrating CD8+ donor T cells (CD45.1- CD45.2+, right) at endpoint. Transmigrated cells more effectively suppressed tumor growth, accumulated in tumors, and enhanced infiltration of host CD8+ T cells (CD45.1+ CD45.2-).(n = 3 mice per group, mean ± SEM; 2 mice in the control group died before day 14, leaving n = 1 for final tumor measurement; *p < 0.05, ****p < 0.0001, one-way ANOVA with Tukey’s multiple comparison test) **(I)** Quantification of donor CD8+ T cells in tumor-draining lymph node (TdLN), non-draining LN (NdLN), and spleen. Transmigrated cells exhibited increased trafficking to secondary lymphoid organs compared to parent cells. (n = 3-4 mice per group; mean ± SEM; **p < 0.01, ***p < 0.001, unpaired t-test)

T cells recovered after transmigration displayed a more polarized morphology and larger size compared to parent cells (Fig. 2E and fig. S12C), features commonly associated with T cell activation (*17*). Moreover, when introduced into three-dimensional (3D) collagen gels of varying density (1.25 vs. 2 mg/mL), transmigrated cells migrated faster than parent cells in 2 mg/ml gels (Fig. 2F), which imposes smaller pore sizes, indicating a mechanically more restrictive microstructure (*18*). Considering the elevated extracellular matrix deposition and resulting mechanically challenging microstructure of solid tumors (*19*), this suggests that transmigrated CD8+ T cells may be better equipped to infiltrate dense tumor tissue and contribute to improved regression of the malignancy *in vivo*.

Whether these functional improvements result in superior tumor control *in vivo* was assessed. Following peritumoral injection into established B16F10-OVA tumors (Fig. 2G), mice that received transmigrated CD8^+^ T cells displayed significantly delayed tumor growth than both untreated and parent cell-treated mice (Fig. 2H). At end point, harvested tumors from mice receiving transmigrated cells not only had more donor (CD45.2^+^) T cells but also host (CD45.1^+^) T cells, suggesting secondary tumor modulation effects such as broader immune activation in addition to the direct elimination of cancer cells. Furthermore, transmigrated cells accumulated more than parent cells in secondary lymphoid organs such as tumor draining lymph nodes and the spleen (Fig. 2I), indicating enhanced systemic trafficking and potential for long-term surveillance. In line with this, phenotypic analysis of donor CD8+ T cells recovered at day 14 (day 7 post-transfer) showed that transmigrated donor cells maintained a lower PD-1+ fraction with higher Ki-67 and GzmB expression, most prominently in the tumor-draining lymph node (fig. S9). These results suggest that T-Chip processing may have effects that persist beyond the immediate post-transfer window.

Altogether, these data demonstrate that physical transmigration transforms CD8^+^ T cells toward a highly migratory phenotype with enhanced cytotoxicity and proliferative capacity, enabling improved tumor infiltration and control. These phenotypic and functional changes were furthermore induced within minutes of transmigration and occurred without genetic engineering, cytokine reprogramming, or pharmacological intervention. Together with the recovery of highly viable transmigrated cells with high expansion capabilities, these findings suggest T-Chip as a platform to interrogate the biology of this T cells response as well as a rapid, reversible, and label-free conditioning strategy to improve the manufacturing of highly potent adoptive T-cell therapies.

### Surface PD-1 remodeling during transmigration is induced rather than selection-driven

To assess whether passive mechanical deformation, rather than an active biological response, is sufficient to trigger acute and selective loss of PD-1 in CD8^+^ T cells, lymphocytes were physically pushed through 3 µm pores using a syringe pump to simulate their passive deformation under convective filtration (Fig. 3A). Under such conditions, PD-1 expression remained unchanged (Fig. 3A, fig. S10) despite extremely high shear stress (>144 dyne/cm^2^), consistent with previous observations (*20*). Intrinsic active transmigration–not passive mechanical deformation–is thus required to trigger acute and selective loss of surface PD-1 in CD8^+^ T cells.

**Figure 3.**
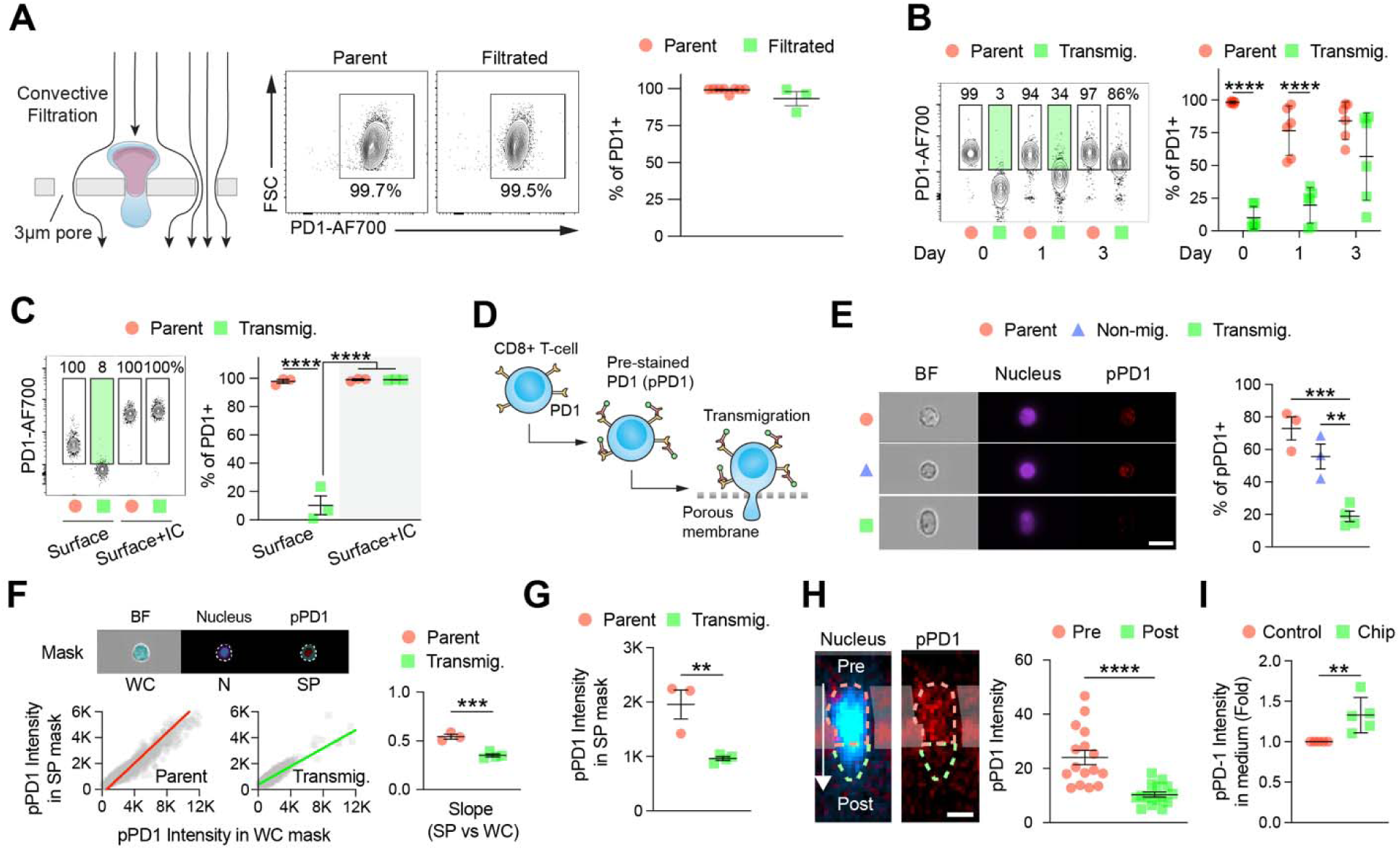
Transmigration triggers acute and selective loss of surface PD-1 in CD8+ T cells. **(A)** Schematic illustration (left), representative flow cytometry plots (top right), and summary (bottom right) of surface PD-1 expression of CD8+ T cells physically pushed through 3 µm pores under high shear stress (144 dyne/cm²) showing no significant reduction compared to parent cells. **(B)** Representative flow cytometry plots (left) and summary (right) of surface PD-1 expression at day 0, 1 and 3 post-transmigration. Green gates highlight the PD-1-low population observed in transmigrated cells at days 0 and 1 post-transmigration. PD-1 expression gradually recovered in transmigrated cells by day 3, while parent cells maintained high PD-1 levels. (n = 3–4 per group; mean ± SEM; ****p < 0.0001, one-way ANOVA with Tukey’s multiple comparison test) **(C)** Representative flow cytometry plots (left) and summary (right) of PD-1 expression on cell surface and whole cell (surface + intracellular) in parent and transmigrated mouse CD8+ T cells. Whole PD-1 staining was performed using the same surface antibody after cell permeabilization to detect both surface and intracellular PD-1. Green gates highlight the reduction in surface PD-1 expression observed in transmigrated cells relative to the parent population. **(D)** Schematic illustration of labeling surface PD-1 on CD8+ T cells (pre-stained surface PD-1, pPD-1) via a fluorescently conjugated anti-PD1 antibody prior to perfusion and transmigration in T-Chip for tracking PD-1 loss. **(E)** Representative imaging flow cytometry images (left) showing pPD-1 (red) and nucleus (magenta) in parent, non-migrated, and transmigrated CD8+ T cells (scale bar = 10 µm). Quantification of pPD1+ cells (right), showing acute pPD-1 loss after transmigration (n = 3-4 chips per group; mean ± SEM; **p < 0.01 ***p < 0.001, one-way ANOVA with Tukey’s multiple comparison test). Full gating strategy for flow cytometry may be found in fig. S12A. **(F)** Analysis of pPD-1 localization using three compartmental masks: whole cell (WC), Nucleus (N), and surface-proximal (SP), calculated as WC minus N. The SP mask represents the membrane-adjacent region where PD-1 is typically localized in imaging flow cytometry images (top left). Representative imaging flow cytometry plots showing pPD1 intensity in SP vs. WC mask for individual cells with linear regression (bottom left). Summary of linear regression slopes (right), indicating a lower surface-to-whole cell pPD-1 intensity slope, thus, surface-dominant PD-1 loss. (n = 3-4 chips per group; mean ± SEM; ***p < 0.001, paired t-test) **(G)** Summary of pPD-1 expression in SPC mask from (G), supporting loss of surface-localized PD-1 after transmigration. (n = 3-4 chips per group; mean ± SEM; **p < 0.01, unpaired t-test) **(H)** Representative cross-sectional confocal images (left) of a transmigrating CD8+ T cell, pre-stained for surface PD-1 (red) and nucleus (blue), within a 3 µm pore (scale bar = 2 µm). Quantification (right) of pPD-1 intensity across pre- and post-transmigration cell bodies. (n = 20 cells per group in 2 experiments; mean ± SEM; ****p < 0.0001, unpaired t-test). White arrow indicates migration direction. **(I)** Fluorometric quantification of pPD-1 intensity in supernatant of parent cells in culture vs. transmigrated cells collected from channels of T-Chip, normalized to parent and expressed as fold change, indicating increased release of pPD-1 by transmigrated T cells. (n = 5 chips per group; mean ± SEM; **p < 0.01, unpaired t-test)

When transmigrated CD8^+^ T cells were recovered and cultured in IL-2 supplemented media, PD-1 levels gradually recovered by day 3, returning to a comparable level to parent cells (Fig. 3B) suggesting a transient modulation mechanism. These findings indicate that surface PD-1 remodeling induced during transmigration is dynamic and reversible. Given prior evidence that T cells can retain intracellular pools of PD-1 that are rapidly mobilized to the surface upon activation (*8, 21, 22*), whether the PD-1 loss observed in transmigrated CD8^+^ T cells was confined to the plasma membrane or reflected a broader reduction in total PD-1 protein expression was interrogated. Surface versus total PD-1 was distinguished by comparing cells stained with and without membrane permeabilization. Despite robust surface PD-1 downregulation, total PD-1 levels remained similar between transmigrated and parent cells (Fig. 3C), implying that intracellular PD-1 depots were preserved.

To directly trace the fate of surface PD-1, CD8^+^ T cells were fluorescently pre-labeled with anti-PD-1 prior to T-Chip processing and imaging flow cytometry was employed to measure spatial PD-1 distributions (pPD1) (Fig. 3, D-G). Total fluorescence intensity of pPD-1 across the whole cell body was significantly reduced in transmigrated cells compared to parent and non-migrated cells (Fig. 3E), implying degradation or extracellular secretion of PD-1 rather than internalization. To further characterize the spatial pattern of surface PD-1 clearance, pPD-1 intensity in the surface-proximal regions and the whole cell at a single-cell level was quantified (Fig. 3F). Both surface proximal and whole cell signals were reduced in transmigrated cells; however, the decrease in surface proximal intensity was significantly greater, resulting in a lower surface proximal to whole cell ratio slope relative to parent cells (Fig. 3, F and G). This single-cell analysis thus highlights a consistent trend of surface-preferential PD-1 clearance, rather than uniform protein loss.

A central concern in interpreting checkpoint loss is whether observed changes reflect active downregulation or selective isolation of pre-existing cells with lower expression levels. This issue, which often complicates interpretation of *in vitro* enrichment studies, is directly addressed in T-Chip using pre-labeled PD-1 imaging. Confocal imaging of T cells during transmigration further supported an induction-based mechanism. T cells pre-stained for PD-1 and imaged mid-transit within micropores via confocal microscopy imaging showed reduced PD-1 signal in cell segments that had passed through the pore, whereas PD-1 was retained on upstream regions (Fig. 3H), implying localized mechanical triggering of PD-1 loss occurring within minutes during transmigration. Real-time imaging of T cells migrating through a horizontal confined microchannel further confirmed rapid and spatially localized loss of pre-labeled PD-1 during transit, consistent with the mid-transit depletion pattern observed in Figure 3I (movie S2 and S3). Elevated pPD-1 signal in the T-Chip supernatant (Fig. 3I) also indicates the release of surface-bound PD-1 into the extracellular space during transmigration, which may also reflect proteolytic shedding, vesicle-associated release, or mechanical detachment of antibody-labeled fragments. These findings collectively demonstrate a selective and rapid loss of surface PD-1 triggered by transmigration within pores, suggestive of post-transcriptional regulation of PD-1 turnover in CD8^+^ T cells.

### Proteasome-mediated degradation underlies PD-1 loss during transmigration

To identify transcriptional signatures of transmigrated T cells, bulk RNA-seq analysis of parent and transmigrated CD8^+^ T cells was performed. Among the differentially expressed genes (DEGs), Pf4, Serpinh1, and Serpinb2 were downregulated, while Tgfbr1, Rprd2, Cirbp, and Nfkbia were upregulated (Fig. 4A), genes all linked to anti-senescence and anti-exhaustion programs (*23–28*). Their coordinated regulation may reflect a shift away from suppressive or arrested transcriptional states. Exploratory analyses of melanoma patient samples in The Cancer Genome Atlas (TCGA) showed that higher expression of Nfkbia, Lilrb4b, C4a and Ccdc141a correlated with longer overall survival (fig. S11). Notably, Rnf19b, which is involved in the ubiquitin-proteasome system, was upregulated (*29*). Gene Ontology enrichment analysis confirmed “proteasome-mediated ubiquitin-dependent protein catabolic process” as the most enriched pathway (Fig. 4B), consistent with recent reports showing that surface PD-1 expression is regulated through E3 ligase–mediated proteasomal degradation in activated T cells (*8*). Exhaustion-associated genes such as Tox and Havcr2 remained unchanged and Lag3 was slightly down-regulated (Fig. 4C). However, Pdcd1 transcript levels were slightly increased, indicating that PD-1 loss is not transcriptionally driven. This finding may reflect compensatory upregulation following acute protein loss, in accordance with gradual recovery of surface PD-1 post-transmigration (Fig. 3B).

**Figure 4.**
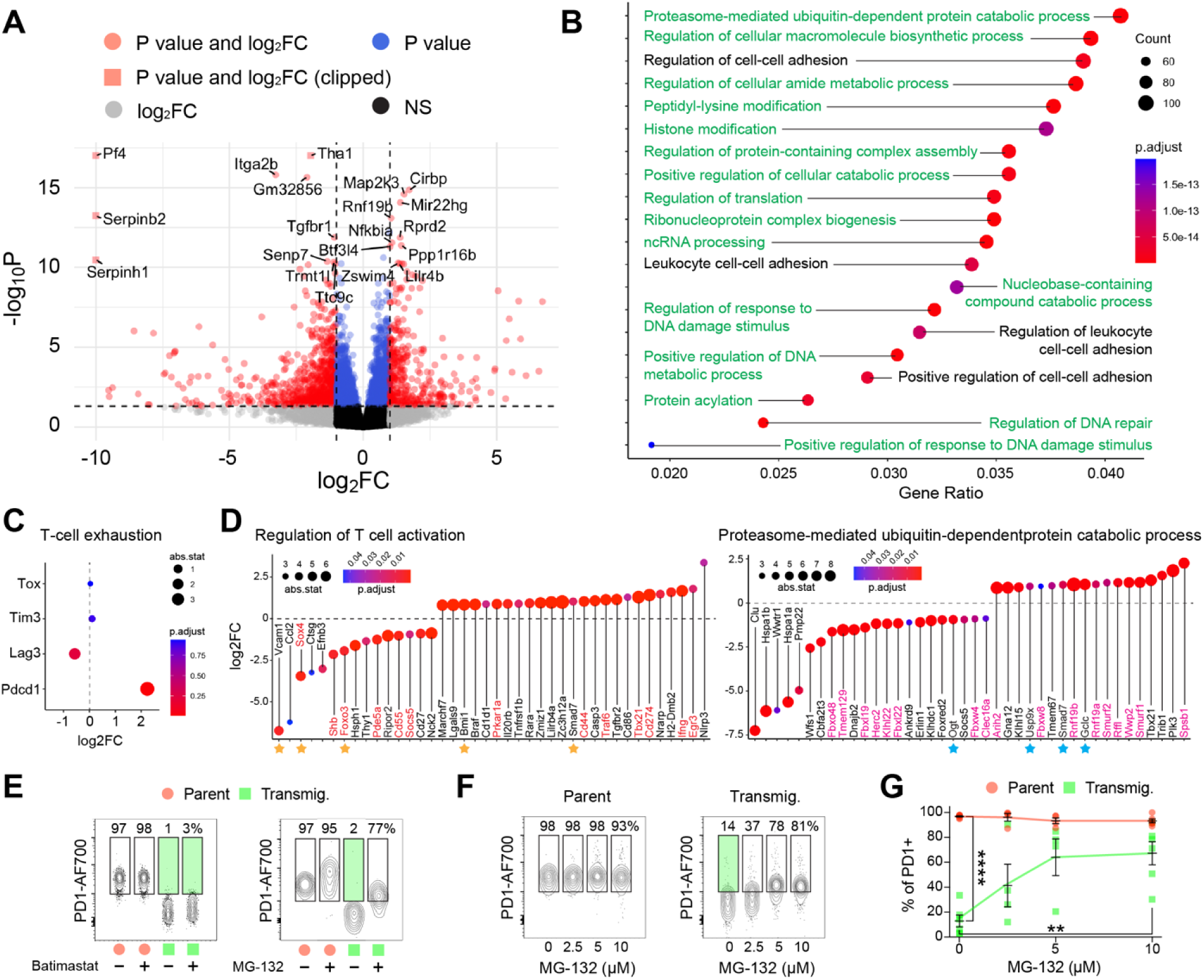
Transmigration promotes proteasome-mediated degradation of PD-1 in CD8+ T cells. **(A)** Volcano plot of differentially expressed genes (DEGs) from bulk RNA-seq comparing parent and transmigrated mouse CD8+ T cells. Dotted lines indicate significance thresholds: adjusted p-value < 0.05 and |log 2 (FC)| > 1 (two-sided Wilcoxon rank-sum test). Data points with extreme values (adjusted p-value < 1e-17 or |log_2_(fold change)| > 10) were clipped to the edge of the plot for visualization. Other genes were categorized: P value group (adjusted p-value < 0.05 and |log 2 (FC)| < 1), log_2_FC group (adjusted p-value < 0.05 and |log 2 (FC)| > 1), and NS (non-significant) group (adjusted p-value > 0.05 and |log 2 (FC)| < 1). **(B)** Dot plot displaying the top 20 enriched Gene Ontology (GO) biological processes. Each dot represents a GO term; dot size indicates the number of genes involved, and color reflects statistical significance (adjusted p-value). The x-axis represents the gene ratio (number of DEGs in the pathway / total number of genes in that pathway). “proteasome-mediated ubiquitin-dependent protein catabolic process” ranked highest by adjusted p-value, indicating increased proteasomal degradation activity. Pathways highlighted in green represent intracellular processing–related pathways. **(C)** Expression of representative T cell exhaustion-related genes (Tox, Tim3/Havcr2, Lag3, Pdcd1) from RNA-seq. Tox and Tim3 were unchanged, Lag3 was down-regulated and Pdcd1 was up-regulated at the mRNA level. **(D)** GO subset enrichment analysis showing top 40 DEGs (ranked by absolute stat value) within “proteasome-mediated ubiquitin-dependent protein catabolic process” and “regulation of T cell activation” categories. Up-regulated genes included E3 ligases (Rnf19a, Rnf19b, Smurf1, Smurf2, Arih2), the E3 adaptor Klhl15, and the deubiquitinase Usp9x, indicating activation of the ubiquitin-proteasome system. Co-occurring up-regulation of Tbx21, Ifng, and Cd44 further suggests that transmigration promotes PD-1 degradation and reprograms CD8+ T cells toward a functionally rejuvenated, effector-like state. Red-labeled genes indicate up- or downregulation reported to enhance effector T cell phenotypes. Yellow star-labeled genes indicate up- or downregulation reported to be associated with anti-exhaustion or anti-senescence. Magenta-labeled genes represent E3 ligase genes. Cyan star-labeled genes indicate up- or downregulation reported to be associated with PD-1 down-regulation. **(E)** Representative surface PD-1 expression in transmigrated CD8+ T cells treated overnight with 100 µM Batimastat (a protease inhibitor) or 10µM MG-132 (a proteasome inhibitor). MG-132, but not Batimastat, prevented PD-1 loss in transmigrated cells, indicating that PD-1 loss is proteasome-dependent. Green gates highlight the PD-1-low population observed in transmigrated cells. **(F)** Representative expression of surface PD-1 expression in (left) parent and (right) transmigrated CD8+ T cells, which were pre-treated overnight with increasing concentrations of MG-132 (0, 2.5, 5, 10 μM) prior to T-Chip processing. Green gates highlight the PD-1-low population observed in transmigrated cells. **(G)** Summary of PD-1+ cells from (G). Surface PD-1 expression on transmigrated cells was recovered in an MG-132 dose-dependent manner, while parent cells remained unaffected. (n = 4-6 chips per group; mean ± SEM; **p < 0.01, ****p < 0.0001, one-way ANOVA with Tukey’s multiple comparisons test)

To further examine other functional categories, differentially expressed genes enriched within “proteasome-mediated ubiquitin-dependent protein catabolic process” and “regulation of T cell activation” were specifically investigated (Fig. 4D). In “regulation of T cell activation” category, transmigrated CD8^+^ T cells exhibited transcriptional reprogramming associated with enhanced effector function (*30–41*), survival, reduced exhaustion, and increased migratory potential. Tbx21 and Ifng were upregulated, indicating effector differentiation (*38, 39*), as was TRAF6, which promote CTLA-4 degradation via ubiquitination, supporting enhanced effector function (*37*). Expression of CD274 and Egr3, which associated with survival and persistence (*40, 41*), were both increased. Importantly, multiple differentially expressed genes indicated attenuated exhaustion. This is evidenced by upregulation of Smad7 and Bmi1, negative regulators of TGF-β–NF-κB crosstalk (*42*) and senescence (*43*), respectively, and by downregulation of Foxo3, a positive regulator of PD-1 and LAG-3 (*44*), Sox4, a transcription factors enriched in exhaustion-associated programs (*45*), and Vcam1, associated with tissue retention in exhausted CD8^+^ T cells (*46*). Together, these changes indicate a potential reversal of exhaustion features. Reduced Ripor2, a negative regulator of RhoA mediated migration (*47*), also supports the enhanced migration in transmigrated cells (Fig. 2F).

Notably, within “proteasome-mediated ubiquitin-dependent protein catabolic process” pathway, several F-box family members (Fbxl19, Fbxl22, and Fbxo48) were downregulated – contrasting with prior findings where E3 ligases such as FBXO38 mediated surface PD-1 degradation (*8*). Despite this result, we observed robust loss of surface PD-1 in transmigrated cells. These findings suggest that mechanical stimuli encountered during transmigration may activate an alternative degradation pathway. Indeed, Gclc, Smad7, and Usp9x were upregulated and Ogt was downregulated, all previously linked to PD-1 turnover, further supporting a transcriptional environment conducive to PD-1 degradation (*42, 48–50*). In addition, multiple E3 ligases including Rnf19a/b, Smurf1/2, and Arih2 were transcriptionally upregulated, and may functionally substitute in targeting PD-1 for degradation, suggesting that mechanical signals in T cell transmigration activates a distinct proteolytic program to downregulate immune checkpoints like PD-1.

Surface PD-1 levels are known to be regulated by various post-translational mechanisms such as shedding by metalloproteinases and ubiquitin-proteasome–mediated degradation (*6, 7*). In light of the transcriptomic enrichment for proteasome-related genes and prior evidence implicating E3 ligase-mediated PD-1 turnover (*8*), we asked whether blocking proteasome activity could rescue surface PD-1 levels. When CD8^+^ T cells were treated with a proteasome inhibitor (MG-132) prior to T-Chip processing, surface PD-1 expression in transmigrated cells was preserved (Fig. 4E), with near-complete rescue at 10 µM (Fig. 4, F and G). In contrast, Batimastat, a broad-spectrum metalloproteinase inhibitor, had no effect, which indicates that intracellular proteasome activity, rather than extracellular shedding, is the main driver of PD-1 loss during transmigration. Together, these data reveal that physical transmigration primes T cells with a transcriptional program favoring PD-1 clearance through proteasome-mediated degradation while also suppressing senescence and promoting pathways of effector function, revealing a novel layer of immune checkpoint regulation linked to mechanical context.

### Proteasome-dependent regulation of surface PD-1 is conserved in human CD8^+^ T cells

Expanded human CD8^+^ T cells were perfused through T-Chips comprised of either 3 µm or 5 µm pores to interrogate whether transmigration-induced PD-1 degradation is conserved in human CD8⁺ T cells. Transmigrated cells showed significantly reduced PD-1, CXCR3, and CD45RA compared to parent populations, with no difference between pore sizes (Fig. 5, A and B). Memory phenotyping revealed a shift toward central memory subsets, with fewer terminally differentiated effector memory cells, contrasting with the effector memory enrichment observed in murine T cells (Fig. 5, C and D). This suggests that migratory stimuli may influence memory subset distribution in a species- and tissue-specific manner. Importantly, PD-1 loss was observed across all memory subsets (Fig. 5D), indicating that transmigration directly remodels PD-1 expression rather than merely selectively depleting specific subsets. Like murine T cells, transmigrated human CD8^+^ T cells migrated faster than parent cells in 3D collagen gels (Fig. 5E) although no morphology polarization was observed (fig. S12D). Additionally, reduced PD-1 expression following transmigration was also observed in conventional Transwell assays (fig. S13), demonstrating that this phenomenon is not unique to the microfluidic platform.

**Figure 5.**
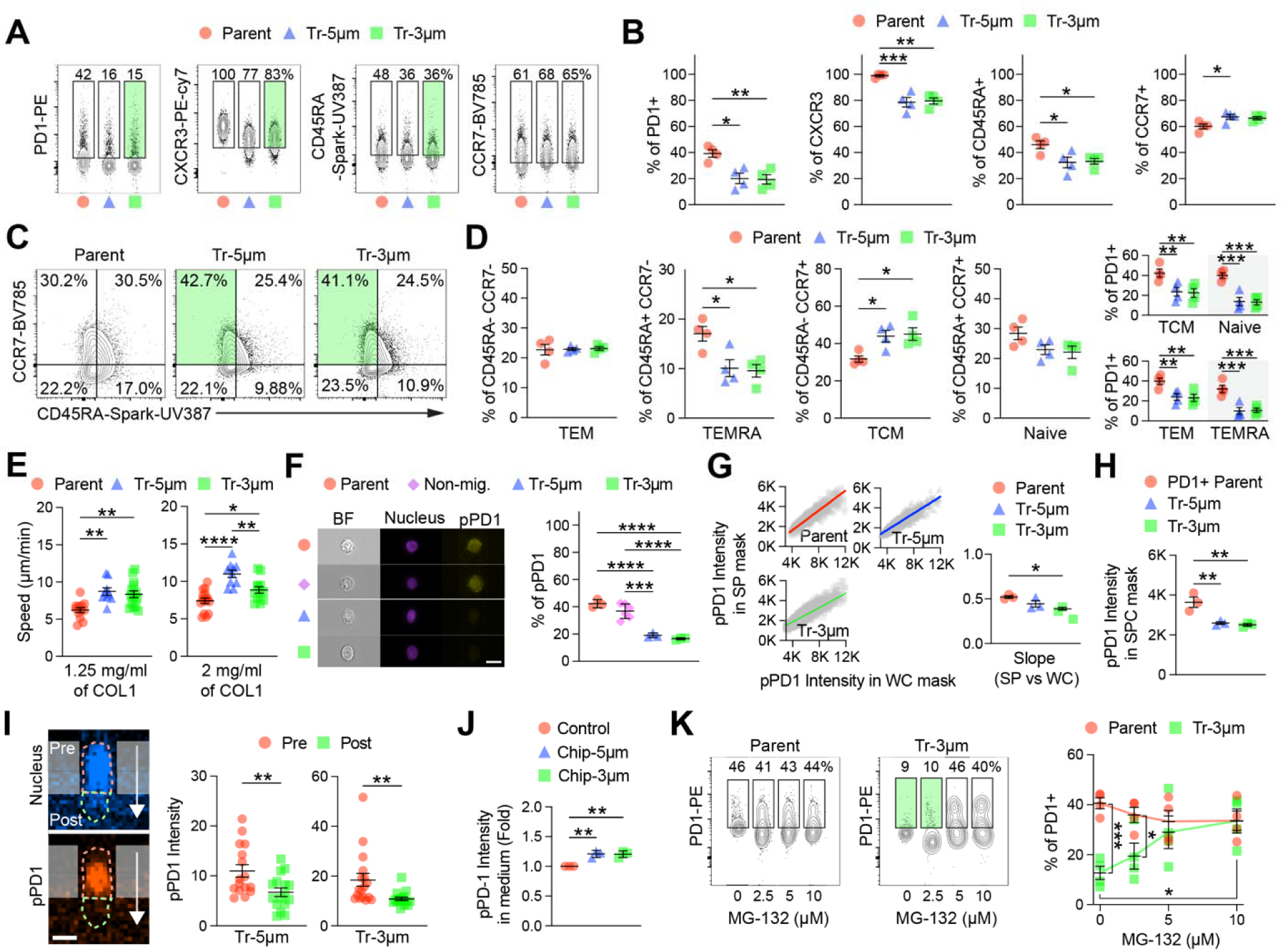
Proteasome-dependent PD-1 degradation is recapitulated in human CD8+ T cells during transmigration in T-Chip. **(A)** Representative flow cytometry plots for expression of surface PD-1, CXCR3, CD45RA, and CCR7 in parent human CD8+ T cells (isolated from peripheral blood mononuclear cells), and transmigrated cells processed through 5 µm (Tr-5µm) or 3 µm (Tr-3µM) pores in the T-Chip. Green gates highlight marker populations that were either enriched or reduced in transmigrated cells relative to the parent population. **(B)** Summary of PD-1+, CXCR3+, CD45RA+, and CCR7+ cells from (A). Transmigrated cells showed reduced PD-1, CXCR3, and CD45RA expression (n = 4 chips per group; mean ± SEM; *p < 0.05, **p < 0.01, ***p < 0.001, one-way ANOVA with Tukey’s multiple comparisons test) **(C)** Representative flow cytometry plots for CD45RA vs. CCR7 expression used for naïve/memory subset classification. Green gates highlight the CD45RA⁻CCR7⁺ central memory population enriched following transmigration. **(D)** Quantification of T cell subset frequencies (left) from (C): naïve (CD45RA+CCR7+), central memory (TCM; CD45RA-CCR7+), effector memory (TEM; CD45RA-CCR7-), and terminally differentiated effector memory (TEMRA; CD45RA+CCR7-). Transmigration decreased TEMRA and increased TCM proportions, indicating a shift toward central memory phenotype. PD-1 expression across different subsets (right). PD-1 down-regulation occurred uniformly across all subsets, suggesting subset agnostic PD-1 reduction. (n = 4 chips per group; mean ± SEM; *p < 0.05, **p < 0.01, ***p < 0.001, one-way ANOVA with Tukey’s multiple comparisons test) **(E)** Migration speed of parent and transmigrated T cells in 3D collagen matrices (1.25 or 2 mg/ml), indicating significantly higher motility of transmigrated T cells (n = 9-15 cells per group in 1-4 experiments; mean ± SEM; **p < 0.01, ****p < 0.0001, one-way ANOVA with Tukey’s multiple comparisons test) **(F)** Representative imaging flow cytometry images (left) showing surface pre-stained PD-1 (pPD1, yellow) and nucleus (magenta) in parent, non-migrated, Tr-5µm, and Tr-3µm cells (scale bar = 10 µm). Quantification of pPD1+ cells (right), indicating acute loss of surface PD-1 following transmigration. (n = 3–6 chips per group; mean ± SEM; ***p < 0.001, ****p < 0.0001, one-way ANOVA with Tukey’s multiple comparisons test) Full gating strategy for flow cytometry may be found in fig. S12B. **(G)** Representative imaging flow cytometry plots (left) showing pPD1 intensity in SPC vs TC mask for individual cells, with linear regression. Quantification of linear regression slopes (right). The reduced slope in Tr-3μm cells indicates preferential loss of surface-localized PD-1. (n = 3 chips per group; mean ± SEM; *p < 0.05, one-way ANOVA with Tukey’s multiple comparisons test) **(H)** Summary of pPD-1 expression remained on cell surface from (G), showing significantly reduced signal in transmigrated cells compared to controls. (n = 3 chips per group; mean ± SEM; **p < 0.01, one-way ANOVA with Tukey’s multiple comparisons test) **(I)** Representative cross-sectional confocal images (left) of a transmigrating human CD8+ T cell, pre-stained for surface PD-1 (orange) and nucleus (blue), within a 3 µm pore (scale bar = 2 µm). Quantification of pPD1 intensity (right) in pre- and post-transmigration segments of the cell body across 3 and 5 µm pores (n = 16 cells from 2 experiments; mean ± SEM; **p < 0.01, unpaired t-test). White arrows indicate migration direction. **(J)** Relative pPD1 signal in culture supernatants from control (parent cells) and T-Chip (transmigrated cells, 5 or 3 µm pores) samples. (n = 3 chips per group; mean ± SEM; **p < 0.01, one-way ANOVA with Tukey’s multiple comparisons test) **(K)** Representative surface PD-1 expression (left) and summary of PD-1+ cell frequency (right) in parent and Tr-3μm cells, which were pre-treated overnight with increasing concentrations of MG-132 (0, 2.5, 5, 10 µM) prior to T-Chip processing. Green gates highlight the PD-1-low population observed in transmigrated cells. MG-132 dose-dependently recovered PD-1 expression in Tr-3µm cells confirming proteasome-dependent PD-1 degradation in human CD8+ T cells (n = 4 chips per group; mean ± SEM; *p < 0.05, ***p < 0.001, two-way ANOVA with Tukey’s multiple comparisons test within-group comparison and Šídák’s multiple comparisons test for between-group comparison)

In pPD-1 tracking, human CD8^+^ T cells exhibited robust PD-1 loss at the cell surface (Fig. 5, F-H). High-resolution confocal microscopy analysis also showed significant preferential clearance in cell regions that had passed through micropores (Fig. 5I), consistent with localized mechanical triggering of PD-1 clearance shown in murine CD8^+^ T cells. pPD-1 intensity was also increased in supernatants collected from T-Chips in comparison to culture medium of parent cells (Fig. 5J), suggesting partial release of pPD-1. Finally, like mouse cells, pre-treatment with MG-132 inhibited PD-1 loss during transmigration of human CD8^+^ T cells in a dose dependent manner (Fig. 5K). Transmigration engages a mechanism for proteasome-dependent PD-1 downregulation that is conserved human T cells.

### Transmigration attenuates inhibitory marker expression in tumor-infiltrating lymphocytes

Given that confined transmigration acutely reduces surface PD-1 in CD8^+^ T cells through a proteasome-dependent mechanism, we next asked whether other inhibitory receptors exhibit similar mechanosensitive dynamics. To this end, short-term adoptive transfer experiments using OT-I CD8^+^ T cells were revisited (Fig. 1L). Notably, Tim-3 expression was markedly reduced in tumor-infiltrating donor cells within 2 h post-injection (fig. S14A) and remained suppressed at 24 h, unlike PD-1, which had largely recovered. Tim-3 remained unchanged in donor T cells recovered from blood (fig. S14B), suggesting that its downregulation is also triggered by tumor entry. Prior studies have shown that Tim-3 can undergo proteasome-mediated degradation in activated T cells (*51*).

We hypothesized that this mechanism could be harnessed to remodel exhaustion-associated phenotypes in therapeutic T cells such as tumor-infiltrating lymphocytes (TIL). To test this idea, expanded TIL from B16F10-OVA tumors treated with high-dose IL-2 and processed them in the T-Chip with 3 µm pores (Fig. 6A). Transmigrated CD8^+^ TILs lost both PD-1 and Tim-3 expression, along with a decreased PD-1^+^Tim-3^+^ cell frequencies and elevated PD-1^−^Tim-3^−^subsets (Fig. 6, B and C). This phenotypic shift was accompanied by more effector memory cells and fewer CD44^−^CD62L^−^ populations (Fig. 6D), consistent with our earlier observations in expanded murine CD8^+^ T cells. Confined transmigration also downregulated surface PD-1 and Tim-3 expression in *ex vivo* expanded human TIL, mirroring effects observed in murine TIL. Following transmigration in T-Chip, human TILs exhibited reduced expression of both markers, along with a shift away from the PD-1^+^Tim-3^+^ population toward the double-negative phenotype (Fig. 6, E–G). Memory subset analysis revealed enrichment of effector memory populations and decreased central memory fractions (Fig. 6H), though donor variability limited statistical significance.

**Figure 6.**
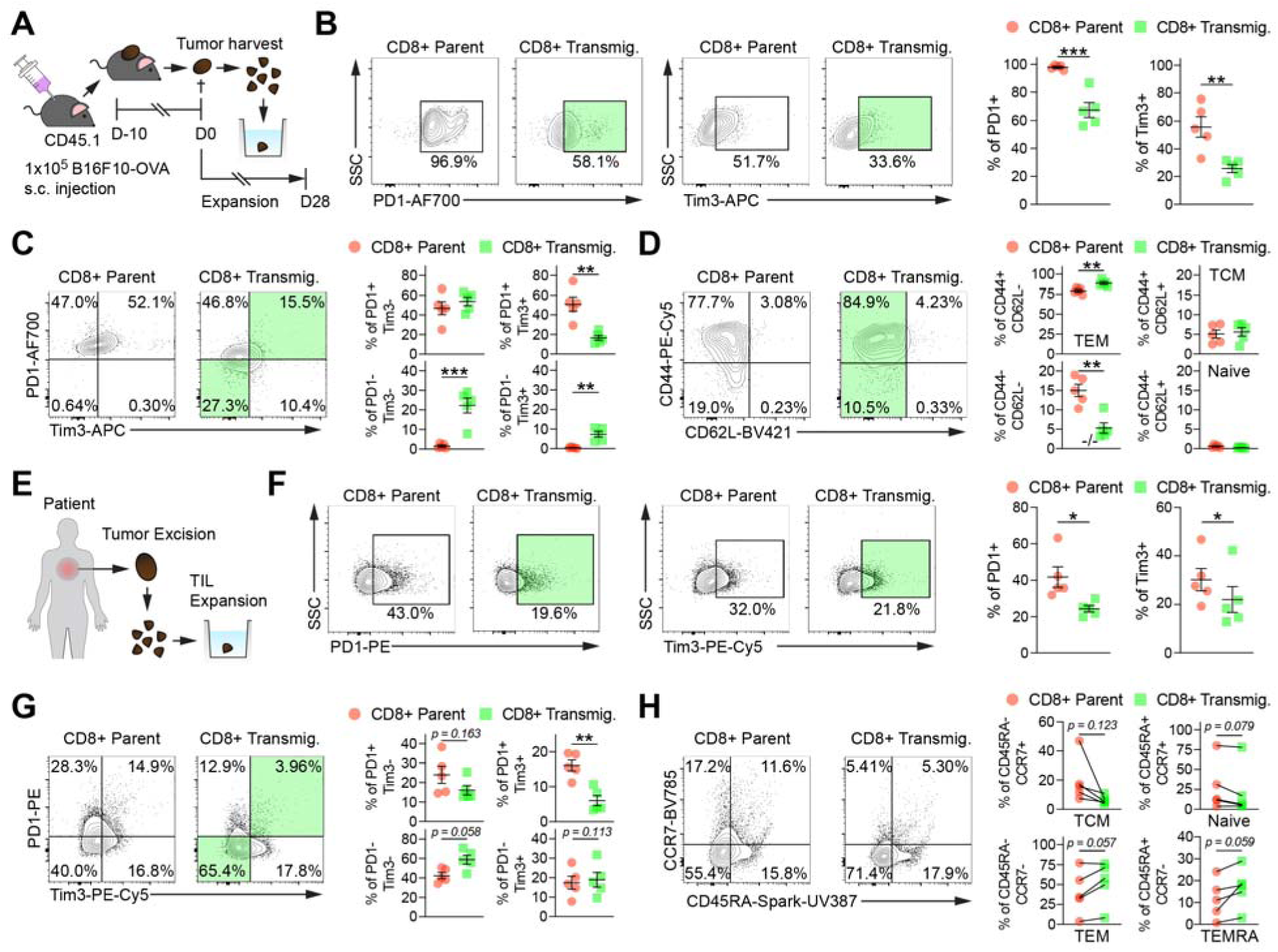
Transmigration-induced PD-1 down-regulation and phenotypic remodeling are reproduced in tumor-infiltrating CD8+ T cells. **(A)** Experimental design of mouse tumor infiltrating lymphocyte (TIL) experiment. Mouse TIL were isolated from B16F10-OVA tumors harvested 10 days after subcutaneous implantation (1×105 cells), expanded ex vivo with high-dose IL-2 (6,000 IU/mL), processed through the TChip (3 μm pores) on day 28 post-expansion and analyzed. **(B)** Experimental design of mouse tumor infiltrating lymphocyte (TIL) experiment. Mouse TILs were isolated from B16F10-OVA tumors harvested 10 days after subcutaneous implantation (1×10^5^ cells), expanded ex vivo with high-dose IL-2 (6,000 IU/mL), processed through the T-Chip (3 µm pores) on day 28 post-expansion and analyzed. **(C)** Representative flow cytometry plots (left) and summary of PD-1 and Tim-3 expression in expanded mouse CD8+ parent and transmigrated TILs. Green gates highlight cell populations with reduced PD-1 or Tim-3 expression following transmigration. Transmigration significantly reduced the expression of both PD-1 and Tim-3. (n = 5 chips per group from 5 independent tumor fragment pools; mean ± SEM; **p < 0.01, ***p < 0.001, unpaired t-test) Full gating strategy for flow cytometry may be found in fig. S15A. **(D)** Representative flow cytometry plots (left) and quantification (right) of PD-1 vs. Tim-3 expression in parent and transmigrated mouse CD8+ TILs. Green gates highlight the increased PD-1-Tim-3- and decreased PD-1+Tim-3+ populations observed following transmigration. Transmigrated cells exhibited increased PD-1-Tim-3- and decreased PD-1+Tim-3+ populations relative to parent cells. (n = 5 chips per group from 5 independent tumor fragment pools; mean ± SEM; **p < 0.01, ***p < 0.001, unpaired t-test) **(E)** Representative CD44 vs. CD62L plots (left) in parent and transmigrated mouse CD8+ TILs and quantification (right) of T cell subsets: double-negative (-/-), naïve (CD44-CD62L+), effector memory (TEM; CD44+CD62L-), and central memory (TCM; CD44+CD62L+) populations. Green gates highlight the TEM population enriched following transmigration. Transmigration increased TEM and reduced -/- populations. (n = 5 chips per group from 5 independent tumor fragment pools; mean ± SEM; **p < 0.01, unpaired t-test) **(F)** Experimental design of human TIL experiment. Human TILs expanded from surgically resected tumors with high-dose IL-2 (6,000 IU/mL) were processed through T-Chip (3 µm pores) and analyzed. **(G)** Representative flow cytometry plots (left) and summary (right) of PD-1 and Tim-3 expressions in parent and transmigrated human CD8+ TIL. Green gates highlight cell populations with reduced PD-1 or Tim-3 expression following transmigration. Transmigration significantly reduced the expression of both PD-1 and Tim-3 in human CD8+ TILs. (n = 5 chips per group from 5 independent tumor fragments; mean ± SEM; *p < 0.05, paired t-test) Full gating strategy for flow cytometry may be found in fig. S15B. **(H)** Representative flow cytometry plots (left) and quantification (right) of PD-1 vs. Tim-3 expression in parent and transmigrated human CD8+ TILs. Green gates highlight the increased PD-1-Tim-3- and decreased PD-1+Tim-3+ populations observed following transmigration. (n = 5 chips per group from 5 independent tumor fragments; mean ± SEM; *p < 0.05, **p < 0.01, unpaired t-test) **(I)** Representative CD45RA vs. CCR7 plots (left) and paired donor analysis (right) for memory subsets in parent and transmigrated human CD8+ TILs. Transmigration reduced TCM (CD45RA-CCR7+) and modestly increased TEM (CD45RA-CCR7-) and TEMRA (CD45RA+CCR7-) populations. Although consistent trends were observed across donors, inter-donor heterogeneity limited statistical significance. (n = 5 chips per group from 5 independent tumor fragments; mean ± SEM; p-values from paired t-test)

Transmigration through the T-Chip furthermore enriched for viable TIL in both murine and human samples (fig. S15, C and D), suggesting that this process not only reprograms exhaustion markers but also serves as a functional purification step that selects for live, responsive T cells. Given the critical importance of viability in adoptive T cell therapies, this enrichment may contribute to improved persistence and antitumor efficacy *in vivo*. We further examined whether this phenotype extends to engineered T cells by applying the same approach to human CAR T cells. Transmigrated CAR T cells displayed robust loss of PD-1 while maintaining CAR expression across a range of CAR expression levels (fig. S16). Together, these findings suggest that confined migration acts as a biophysical cue that tunes exhaustion phenotypes and functionality of both endogenous and engineered T cells.

## Discussion

Our work identifies mechanical confinement during transmigration as an unrecognized checkpoint regulator, revealing that biophysical cues can acutely reprogram surface receptor expression and functional states. T cell immune checkpoint expression is widely regarded a consequence of chronic antigen or inflammatory cytokine stimulation, reinforcing the paradigm that T cell dysfunction is orchestrated primarily by molecular cues. This dogma is now challenged by observations of mechanical checkpoints impinged by biophysical interactions with cancer cells and stiff extracellular matrix of tumors inducing regulation of surface checkpoint inhibitors. While direct interactions within tumors are crucial for T cell function, the role of trans-endothelial migration, which is performed routinely by T cells and requiring transient migration in an extremely confined microenvironment, remains largely unexplored.

Our results show that confined migration triggers rapid and selective downregulation of surface PD-1 in CD8^+^ T cells via a proteasome-dependent mechanism. We propose that mechanical confinement, alongside well-characterized antigenic and cytokine-mediated inputs, actively shapes checkpoint expression and inhibitory programs in T cells. This acute downregulation was accompanied by transcriptional and functional remodeling. These shifts imply that T cells may transiently reduce exhaustion-associated phenotypes and enhance effector readiness in response to mechanical cues encountered upon entering inflamed tissues. Importantly, this interpretation is supported by acute PD-1 downregulation observed following intravenous but not intratumoral transfer, despite both populations being analyzed within the same tumor microenvironment. Because intratumoral injection bypasses vascular trafficking and trans-endothelial migration, these findings support tissue entry itself, rather than exposure to tumor-derived factors, is a primary driver of the early checkpoint remodeling observed in infiltrating T cells. Trans-endothelial migration thus likely serves as a “reset point” during tissue entry—briefly removing surface checkpoints, priming motility and proteolytic programs, and enhancing environmental responsiveness. These findings reinforce the emerging view that surface checkpoint expression and inhibitory tone can be dynamically remodeled in response to defined cues (*52, 53*), including mechanical ones. Moreover, our results raise the intriguing possibility that PD-1, classically viewed as a marker of antigen experience, may also provide insight into the timing of tissue integration or residency, offering a new perspective on its biological role. Tissue entry may also constitute an additional, Fcγ receptor-independent mechanism capable of reducing checkpoint receptor mAb occupancy and thus the extent of therapeutic checkpoint inhibition achieved in vivo.

T cell exhaustion or dysfunction, associated with checkpoint expression in chronically inflamed microenvironments, such as solid tumors, remains one of the most pressing bottlenecks in cancer immunotherapy (*54*). Our findings that confined migration downregulates both PD-1 and Tim3 positions transmigration as a novel axis for therapeutic checkpoint modulation. Notably, engineered *ex vivo* transmigration could be harnessed to reinvigorate checkpoint-high tumor-infiltrating T cells before adoptive therapy. Unlike genetic or pharmacological interventions, this approach is rapid, reversible, label-free, and highly scalable potentially enabling seamless integration into TIL and CAR T manufacturing pipelines. This concept is particularly timely considering the recent FDA approval of lifileucel (*55*), the first TIL-based therapy for metastatic melanoma. Moreover, this T-Chip–mediated transmigration may function as a “mechanical adjuvant” compatible with engineered cell therapy platforms, including CRISPR-edited, T cell receptor-engineered, or CAR T cells. In our study, CAR T cells retained transgene expression following transmigration while reducing PD-1, supporting the integration of this approach as a transient, label-free checkpoint attenuation step. Beyond final-stage processing, confined migration could also be incorporated earlier in *ex vivo* T cell expansion workflows to mitigate early exhaustion (*56*) and support more robust proliferation.

While our findings establish confined migration as a trigger for proteasome-dependent checkpoint clearance, several mechanistic questions remain: (i) defining upstream mechanosensors and cytoskeletal programs that engage ubiquitin-proteasome pathways, (ii) determining how transient versus chronic confinement shape T cell states, and (iii) resolving subset heterogeneity with single-cell or spatial approaches. Beyond trans-endothelial migration, traversing tumor microenvironments also requires migration in confined porous spaces formed within stiff and dense extracellular matrix fibers (*5, 57*). This mechanically challenging tissue microenvironment is in general associated with T cell arrest, exclusion, and exhaustion (*5, 57, 58*). This likely reflects the temporal nature of two distinct physiological processes, where transmigration exposes cells to a confined environment acutely but transiently and, on the other hand, the tumor microenvironment imposes chronic mechanical confinement with added immunosuppressive interactions. It remains to be determined whether biophysical parameters like pore size, stiffness, or dwell time can be tuned to program distinct T cell fate. Clarifying these mechanisms, a program T-Chip is well positioned to enable, will not only deepen understanding of T cell mechanobiology but also guide the rational design of mechanical pre-conditioning strategies for cellular immunotherapies.

Overall, our results reveal mechanical interactions can reverse dysfunctional state of T cells and regulate their surface protein expression, which is functionally employed in living organisms in the form of trans-endothelial migration. Not only can these physiological concepts be translated into *ex vivo* mechanical modulation of T cells via T-Chip platform as a next-generation reprogramming strategy for adoptive cell therapies, but they also illustrate the pervasive role biophysical forces play in modulating immunity.

## Materials and Methods

### Mice and tumor model

C57BL/6 (CD45.2) and congenic B6 CD45.1 female mice were purchased at six weeks of age from the Jackson Laboratory. OT-1 transgenic mice were bred in-house from breeding pairs obtained from the Jackson Laboratory. All mice were maintained under specific pathogen–free conditions in the Georgia Institute of Technology Animal Facility. All protocols were approved by the Institutional Animal Care and Use Committee.

B16F10-OVA or parental B16F10 melanoma cells were cultured in Dulbecco’s modified Eagle’s medium supplemented with 10% fetal bovine serum, 1% penicillin/streptomycin. Cells were passaged at ∼80% confluence and maintained at 37 °C with 5% CO2 in a humidified incubator. Cells were used for implantation at ∼80% confluence. For tumor implantation, 1×10L B16F10-OVA melanoma cells were injected subcutaneously into the right flank of CD45.1⁺ mice 7 days prior to adoptive transfer of T cells. To evaluate therapeutic effects, tumor size was measured with calipers in three dimensions and reported as an ellipsoidal volume.

### Mouse T cell expansion and adoptive transfer

Spleens were harvested from C57BL/6 or OT-I mice following euthanasia. Cells were prepared by passing spleens through a sterile 70-µm cell strainer and lysing red blood cells with RBC lysis buffer (Sigma-Aldrich). CD8⁺ T cells were isolated using a MojoSort Mouse CD8 T Cell Isolation Kit (BioLegend) according to the manufacturer’s instructions. Purified cells were activated using Dynabeads Mouse T-Activator CD3/CD28 (Thermo Fisher Scientific) at a 1:1 bead-to-cell ratio and cultured in RPMI 1640 medium supplemented with 10% fetal bovine serum, 1% penicillin–streptomycin, 1% HEPES, 1% non-essential amino acids, 1% sodium pyruvate, 0.05 mM 2-mercaptoethanol, and 100 U/ml recombinant IL-2. Cells were monitored daily, and cells were split every 2 days after day 3. Beads were magnetically removed on day 5.

For short term homing experiments, CD8⁺ OT-I T cells expanded for 10 or 11 days were intravenously injected into tumor-bearing CD45.1⁺ mice at a dose of 7.5×10L cells per mouse. Tumors were harvested at 2- and 24-hours post-transfer for analysis.

For intratumoral versus intravenous delivery comparison, CD45.1⁺ recipient mice bearing parental B16F10 tumors received 0.5×10L or 2.0×10L expanded CD45.2⁺ CD8⁺ T cells either by intravenous injection or by intratumoral injection respectively. Donor T cells (CD45.1⁻ CD45.2⁺ CD8⁺) were recovered from tumors at 2 h post-transfer and analyzed.

For pre-labeled anti–PD-1 adoptive transfer with FcγR blockade, expanded OT-I CD45.2⁺ CD8⁺ T cells were pre-labeled in vitro with fluorescent anti–PD-1 (Clone 29F.1A12) for 1 h prior to adoptive transfer and injected intravenously at a dose of 2.0×10L cells per mouse into CD45.1⁺ B16F10-OVA tumor-bearing recipient mice. Recipient mice received 200 µg of Fcγ receptor (FcγR) blocking antibody (Clone 2.4G2, BioXCell) by intraperitoneal (i.p.) injection 2 h before transfer to block FcγR-mediated antibody stripping. Donor T cells (CD45.1⁻ CD45.2⁺ CD8⁺) were recovered from blood and tumor 2 h post-transfer and analyzed for anti–PD-1 signal by flow cytometry.

For tumor control experiments, expanded CD8+ OT-I T cells were either directly injected as parent cells or injected after undergoing transmigration through T-Chip. Both groups were peritumorally injected into tumor-bearing CD45.1+ mice at a dose of 3×10⁴ cells per mouse. Tumors were harvested 7 days post-transfer. Tumor volume was measured using calipers in three dimensions and calculated as ellipsoidal volume.

### Flow cytometry

For single-cell suspension preparation, harvested lymph nodes, spleens, and tumors were incubated in D-PBS containing 1 mg/mL collagenase D (Sigma-Aldrich) at 37 °C. Lymph nodes and spleens were digested for 1 hour, and tumors for 4 hours. Tissues were then filtered through a 70 μm cell strainer. Blood was collected via retro-orbital bleed, and red blood cells in spleen and blood samples were lysed using RBC lysis buffer (Sigma-Aldrich). All antibodies were obtained from Biolegend unless otherwise specified. Prior to staining, mouse and human cells were blocked with anti-mouse CD16/CD32 (clone 2.4G2, Tonbo Biosciences) or Human TruStain FcX (BioLegend), respectively, for 5 minutes at room temperature. Live/dead staining was performed using Zombie Aqua or Zombie Violet fixable viability dyes (BioLegend), incubated for 15 minutes at room temperature. Antibody cocktails were prepared in flow cytometry buffer (0.1% bovine serum albumin in D-PBS) following manufacturer concentration or preliminary titrations. Surface staining was performed by incubating cells with the antibody cocktail for 25 minutes at 4 °C. For intracellular staining, cells were fixed and permeabilized using the Foxp3/Transcription Factor Staining Buffer Set (eBioscience, Thermo Fisher Scientific), followed by incubation with intracellular antibody cocktails in permeabilization buffer for 25 minutes at 4 °C. Samples were acquired using a Cytek Aurora spectral flow cytometer and analyzed with FlowJo (Tree Star). All antibodies and reagents used for flow cytometry are listed in Table S1.

### T-Chip Fabrication

The microfluidic transmigration-on-a-chip (T-Chip) device was constructed from five stacked layers: a top cover, an upper perfusion channel layer, a microporous membrane (3 μm or 5µm pore size, 6×10L or 2×10L pores/cm²; Sterlitech), a lower chemokine reservoir channel, and a bottom cover (Figure S2). The upper channel layer was fabricated by laser cutting double-sided adhesive (DSA) sheets (50 µm thickness; 3M) into the desired microchannel geometry, aligned and attached onto a polymethyl methacrylate (PMMA) top cover with laser micromachined inlets and outlets. The lower channel layer was also fabricated by laser cutting a 1Lmm-thick PMMA sheet with 50Lμm-thick double-sided adhesive layers attached to both the top and bottom surfaces, resulting in a total channel height of 1.1 mm, and bonded to a flat PMMA bottom cover. The microporous membrane was placed between the upper and lower channel layers, separating the upper and lower channels while enabling diffusion of chemoattractant molecules and transmigration of cells. Inlet and outlet ports were connected to sterile biomedical-grade silicon tubing and sealed to prevent leakage. Assembled T-Chips was sterilized using ethylene oxide gas followed by ultraviolet light prior to use. Computational modeling of fluid velocity and chemokine diffusion in the T-Chip was performed using COMSOL Multiphysics. Detailed simulation parameters and boundary conditions are provided in the Supplementary Methods.

### Transmigration assay

T-Chip was sequentially primed with 70% Ethanol, deionized water, and 1x PBS. To enable T cell rolling and firm adhesion, the upper channel was functionalized with recombinant mouse or human adhesion molecules (R&D Systems) by injecting 1x PBS containing E-selectin (20 µg/mL) and P- selectin (20 µg/mL), VCAM-1 (20 µg/mL), and ICAM-1 (20 µg/mL), and incubating at room temperature for 30 mins. To induce chemokine-guided transmigration, the lower channel was filled with the T cell culture medium supplemented with chemokine cues: CCL2 (30 ng/mL), CCL3 (25 ng/mL), CCL4 (24 ng/mL), CCL5 (100 ng/mL), CXCL9 (300 ng/mL), CXCL10 (300 ng/mL). The chemokine medium was replenished every 30 mins during the assay. Expanded CD8+ T cells were resuspended in the T cell culture medium at 1.0×10^6^ cells/ml (human) or 0.5×10^6^ cells/mL (mouse), and perfused through the upper channel at a wall shear stress of 0.75 dyn/cm^2^ using a syringe pump (Harvard Apparatus) for 3 hours at 37L°C. Following perfusion, non-migrated cells were perfused out and collected, while the cells recovered from the lower channel were collected separately as transmigrated populations. The recovered cells were counted using hemocytometer and processed for downstream analysis or further culture. In selected experiments, adhesion molecules or chemokines were omitted to assess their contribution to transmigration efficiency, defined as % of live cells recovered from the lower channel relative to the number of live cells introduced. To visualize T cell behavior in real-time during the transmigration assay, live-cell imaging was performed on an inverted microscope (Eclipse Ti optical microscope; Nikon) equipped with temperature control. Time-lapse images were captured every 30 s using a 20× objective. Cell rolling, adhesion, and transmigration events were manually annotated using ImageJ. Cell size and shape factor was measured using ImageJ.

For selected experiments, expanded CD8 T cells were pre-treated overnight with Batimastat (100 µM; Sigma-Aldrich) or MG-132 (2.5, 5, or 10 µM; Sigma-Aldrich), which were dissolved in DMSO, in the T cell culture medium. The final concentration of DMSO did not exceed 0.5 % (v/v) in any condition. Control groups were treated with vehicle (DMSO) alone. Following incubation, cells were immediately used in the transmigration assay in the presence of the inhibitors in both the upper and lower channels.

### Transwell transmigration assays

Immortalized mouse lymphatic endothelial cells (LECs), a kind gift from Dr. J. Brandon Dixon, were seeded onto transwell inserts (3 µm or 8 µm pore membranes; Corning) at 5×10^4^ cells/mm^2^ and cultured for 3 days in endothelial basal medium (Sciencell) with 10% FBS to form confluent monolayers. The expanded murine CD8⁺ T cells were added to the upper chamber at 3×10^4^ cells/mm^2^, and chemokine cues were added to the lower chamber. After 4 hours, transmigrated cells were collected from the lower chamber and analyzed by flow cytometry for surface PD-1 expression. Membrane-only inserts (no LECs) were included as controls for each pore size. Live confocal 3D images were performed using a Zeiss LSM900 microscope and analyzed by ImageJ.

### T cell re-stimulation assay

B16F10 OVA cells were cultured to 80% confluence and treated with IFN-γ (5, 10, 20, 40 ng/mL) for 24 h. Based on PD-L1 upregulation saturation (Figure S5), 10 ng/ml IFN-γ was selected for all subsequent experiments. Tumor cells, either untreated or IFN-γ treated, were plated in 96-well plates at 62,500 cells per well and incubated for 2 h prior to T cell co-culture. For intracellular cytokine staining, 62,500 parent or transmigrated T cells were added in the presence of Brefeldin A (5 µg/ml) and incubated for 3 h. All cells were collected and analyzed by flow cytometry. For cytotoxicity assay, parent or transmigrated T cells were labeled with CellTrace Blue for 20 min, washed, and co-cultured with untreated or IFN-γ treated tumor cells (62,500 cells per well) in the presence or absence of anti-PD1 (10 µg/ml). After 24 h, all cells were harvested and analyzed by flow cytometry. Proliferating populations were quantified using FlowJo’s proliferation modeling tool based on CellTrace Blue dilution.

### Collagen migration assay

An assay chamber (4 mm x 4mm x 6 mm height) was fabricated by laser cutting PMMA sheet and bonding it to cover glass using DSA. The fabricated chamber was sterilized with 70 % ethanol and ultraviolet light prior to use. Type 1 collagen (BD) was neutralized by with 10x MEM, 0.5 N NaOH, and deionized water to reach a final concentration (1.25 or 2 mg/ml), pH7.4, and isotonic conditions. The first collagen layer was cast into the chamber and allowed to polymerize in a CO2 incubator. Expanded CD8 T cells were re-suspended in pre-neutralized collagen solution (1.25 or 2 mg/ml) at 1.0×10L cells/ml and layered on top of the first gel. After gelation of the second layer, a third collagen layer was added to embed the cells. The chamber was filled with T cell culture medium and sealed with a top cover glass. Live-cell imaging was performed at 10-second intervals for 15 minutes. Individual cell trajectories were tracked using ImageJ, and migration velocity was quantified by measuring total displacement over time.

### Mechanical compression assay

A circular flow chamber was fabricated from five stacked layers: a top cover, an upper chamber, a microporous membrane (3 μm pore size, Sterlitech), a lower chamber, and a bottom cover (Figure S9). The upper chamber and lower chamber have a height of 1.5 mm. Inlet and outlet ports were connected to sterile biomedical-grade silicon tubing and sealed to prevent leakage. Total 2×10^6^ expanded CD8+ T cells were resuspended in the T cell culture medium at 0.25×10^6^ cells/mL and perfused through the upper chamber at a wall shear stress of 144 and 287 dyn/cm^2^ using a syringe pump (Harvard Apparatus). Following perfusion, the cells recovered from the outlet were collected and analyzed.

### PD-1 tracking experiment

Expanded T cells were blocked with anti-mouse CD16/CD32 or Human TruStain FcX, respectively, for 5 minutes at room temperature and stained with a fluorescently conjugated anti-PD-1 antibody along with hoechst 33342 for 25 minutes at 4 °C. Pre-stained T cells were resuspended in the T cell culture medium at 1.0×10L cells/ml (human) or 0.5×10L cells/mL (mouse), and perfused in T-Chip. Following perfusion, the culture medium used for both the upper and lower channels was collected separately and centrifuged. The culture medium collected from pre-stained parent cells was also prepared as a control. The supernatants were transferred into black-walled 96-well plate for fluorescence analysis using a plate reader. Pre-stained parent, non-migrated cells, and transmigrated cells were fixed using the Foxp3/Transcription Factor Staining Buffer Set. Image flow cytometry was performed using the Amnis ImageStreamX Mk II Imaging Flow Cytometer (Cytek). Data were analyzed with IDEAS software (Image Data Exploration and Analysis Software). After T-Chip disassembly, the porous membranes were carefully retrieved, fixed with the same buffer set, and washed. The membranes were mounted onto cover glasses using VECTASHIELD Vibrance Antifade Mounting Medium (Vector Laboratories) and sealed with an additional cover glass. Confocal 3D images were performed using a Zeiss LSM900 microscope and analyzed by ImageJ.

### Bulk RNA-sequencing and analysis

Total RNA was extracted from parent and transmigrated T cells using the RNeasy Plus Micro Kit (Qiagen), following the manufacturer’s protocol. Quality control and alignment of FASTQ files were performed using the standard workflow at the Emory Integrated Computational Core. Briefly, sequencing was performed on an Illumina NovaSeq 6000 platform. Sequences were quality checked using FastQC, were aligned to the MM39 reference genome using STAR aligner. Gene-level read counts were quantified using HTSeq-count. Differential gene expression analysis was performed using DESeq2. Low count genes (mean count < 5) and outliers (Cook’s distance) were excluded. Significantly differentially expressed genes were defined by adjusted p-value (FDR) < 0.05 and |log2 fold change| > 1. Gene set enrichment analysis (GSEA) was conducted using Reactome and Gene Ontology Biological Process (GO:BP) categories to identify enriched pathways. Visualization was performed in RStudio.

### Human T cell expansion

Human peripheral blood mononuclear cells (PBMC) were isolated from leukopaks obtained from healthy donors under IRB-approved protocols (IRB00100041). PBMC were separated using Ficoll-Paque Premium density gradient centrifugation (Cytiva) following the manufacturer’s instructions. CD8+ T cells were purified by negative selection using the MojoSort Human CD8 T Cell Isolation Kit (BioLegend). Purified cells were activated with Dynabeads Human T-Activator CD3/CD28 (Thermo Fisher Scientific) at a 1:1 bead-to-cell ratio and cultured in the T cell culture medium at 37 °C in a humidified incubator with 5% CO2 for 11 days. Beads were magnetically removed prior to cryopreservation. For T-Chip experiments, cells were thawed and immediately used. Anti-CD19 human CAR T cells were produced as previously reported (*59*). Briefly, PBMC were activated with the Phytohemagglutinin for 2 days, transduced retrovirally for 3 consecutive days, and allowed to rest for 2 days. The CAR T cells were thawed and immediately used for T-Chip experiments.

### Tumor infiltrating lymphocyte expansion

TILs were expanded ex vivo following a previously described protocol (*60*). B16F10-OVA tumors were established in CD45.1+ C57BL/6 mice by subcutaneous injection of 1×10L tumor cells into the right flank. Tumors were harvested 10 days post-implantation, before reaching a maximum size of 10×10 mm². Under sterile conditions, tumors were dissected into ∼2–3 mm² fragments and individually plated in wells of a 24-well plate containing 2 mL of the T-cell culture medium supplemented with 6,000 IU/mL recombinant IL-2. Cultures were maintained undisturbed for the first 5 days. Thereafter, half of the medium was replaced 2–3 times per week. TILs were split as needed to maintain a density of ∼1×10L cells/mL. Tumor fragments were removed either at the first split or depending on the rate of lymphocyte egress. Expanded TILs were collected on day 28, cryopreserved, and used immediately for T-Chip perfusion after thawing.

For Human TIL expansion, human tumor fragments were processed for TIL expansion as previously described (*61*). Deidentified human tumor tissues were collected under Emory University IRB approval (STUDY00001671) from patients undergoing surgical resection. Tumor samples were dissected into fragments approximately 3mm in diameter. Within a 24-well plate, a single fragment per well was cultured for two to five weeks in complete media (RPMI 1640 with L-glutamine with 10% heat-inactivated FBS, 1% penicillin/streptomycin, 1% nonessential amino acids, 1% sodium pyruvate, 0.1% HEPES, and 0.1% β-mercaptoethanol) containing 6000 IU mL^-1^ rhIL-2 (NCI BRB Preclinical Repository) in an incubator at 32**°**C, 5% CO_2_. Egressed and expanded tumor infiltrating lymphocyte (TIL) cultures were cryopreserved until use in this study.

### Statistical analysis

All statistical tests, except for bulk RNA-seq, were performed using Prism 10 software (GraphPad) and are described as follows. Comparisons between two groups were conducted using unpaired two-tailed Student’s t-test unless otherwise noted. For comparisons among multiple groups, one-way ANOVA with Tukey’s multiple comparison test was used. When analyzing two independent variables in multiple groups, two-way ANOVA was used with Tukey’s multiple comparison test or Šídák’s multiple comparisons test. P values less than 0.05 were considered statistically significant. For all figures, *P < 0.05, **P < 0.01, ***P < 0.001, ****P < 0.0001. Data are presented as mean ± SEM unless otherwise indicated. Data are presented as mean ± SEM unless otherwise noted. Individual data points represent biological replicates where applicable, and the number of replicates (n) is indicated in figure legends.

## Acknowledgements

This study was supported in part by the Emory Integrated Computational Core (EICC) (RRID:SCR_023525), which is subsidized by the Emory University School of Medicine and is one of the Emory Integrated Core Facilities. Additional support was provided by the National Center for Georgia Clinical & Translational Science Alliance of the National Institutes of Health under Award Number UL1TR002378. The content is solely the responsibility of the authors and does not necessarily reflect the official views of the National Institutes of Health.

## Author Contributions

S.N.T. and Y.A. conceived the project. Y.A. and J.K. performed all device design and fabrication. All authors designed the experiments. A.S. and T.H.Y. performed image analyses. Y.A., J.K., K.M., S.N.L, and P.A.A. performed *in vivo* experiments. S.R., C.P., H.L., R.F., and M.M.W. provided cells and scientific input on data interpretation. J.K., S.N.L., and M.C.W. performed RNA-seq data analysis and interpretation. The manuscript was written by J.K., Y.A., and S.N.T. All authors discussed the results and edited the manuscript.

## Data and materials availability

The results presented in fig. S11 are in part based upon data generated by the TCGA Research Network (https://www.cancer.gov/tcga). All data are available in the main text or the supplementary materials.

