## Supplementary information for "Confined T cell migration controls programmed cell death 1 expression"

**The PDF file includes:**

Supplementary Methods

Figs. S1 to S16

Tables S1

**Other Supplementary Materials for this manuscript include the following:**

Movies S1 to S3

**Supplementary Methods**

**Additional In vivo short-term trafficking assays**

Short-term adoptive transfer experiments were performed as described in the main Methods for tumor establishment, donor T-cell preparation, tissue harvest, and flow cytometry staining/gating, with the following condition-specific modifications. For OVA versus non-OVA tumor comparison, CD45.1⁺ recipient mice were implanted subcutaneously with either B16F10-OVA or parental B16F10 tumor cells as indicated. Expanded OT-I CD45.2⁺ CD8⁺ T cells were transferred by intravenous injection. Donor CD8⁺ T cells (CD45.1⁻ CD45.2⁺ CD8⁺) were recovered from tumors and tumor-draining lymph nodes (TdLNs) at 2 h and 24 h post-transfer and analyzed for surface PD-1 expression by flow cytometry.

**Computational simulation**

Computational fluid dynamics and mass transport simulations were performed using COMSOL Multiphysics 6.2. The three-dimensional geometry of the T-Chip was reconstructed based on the fabricated device dimensions. For flow simulations, incompressible laminar flow was assumed, and culture medium was modeled as a Newtonian fluid with the properties of water at 37°C. A volumetric flow rate corresponding to the experimental condition (0.75 dyn/cm² wall shear stress) was applied at the inlet, and zero-pressure boundary conditions were assigned at the outlet. Steady-state velocity profiles were calculated throughout the upper channel network. To evaluate chemokine transport, diffusion of CXCL10 from the lower channel was simulated concurrently with laminar flow in the upper channel using the Transport of Diluted Species module. An initial CXCL10 concentration corresponding to experimental conditions was assigned to the lower channel, and transport across the porous membrane was modeled by diffusion. The diffusion coefficient of CXCL10 was set to 1.67×10^-10^ m²/s.

**Transwell transmigration assays**

For human Transwell transmigration assays human lymphatic endothelial cells (LECs; ScienCell) were seeded and cultured on transwell inserts under the same conditions as above to generate confluent monolayers. Expanded human CD8⁺ T cells were added to the upper chamber, and transmigration was performed for 4 h with chemokine cues in the lower chamber. Transmigrated cells were collected from the lower chamber and analyzed by flow cytometry for surface PD-1 expression, with membrane-only inserts included as controls.

**Real-time imaging of PD-1 dynamics during confined microchannel migration**

For real time monitoring of migrating PD-1 pre-stained mouse CD8⁺ T cells, a confined channel (width × height = 5 µm × 5 µm) device was fabricated by casting polydimethylsiloxane (PDMS; 10:1 base-to-curing agent) onto a high-resolution 3D-printed mold produced using a Nanoscribe Photonic Professional GT2 system. PDMS was degassed to remove bubbles and cured at 90°C for at least 3 h. Inlet and outlet ports were created using a biopsy punch. The cured PDMS replica was bonded to a glass substrate by oxygen plasma treatment using a plasma cleaner (Harrick Plasma). Live confocal 3D images were performed using a Zeiss LSM900 microscope and analyzed by ImageJ.

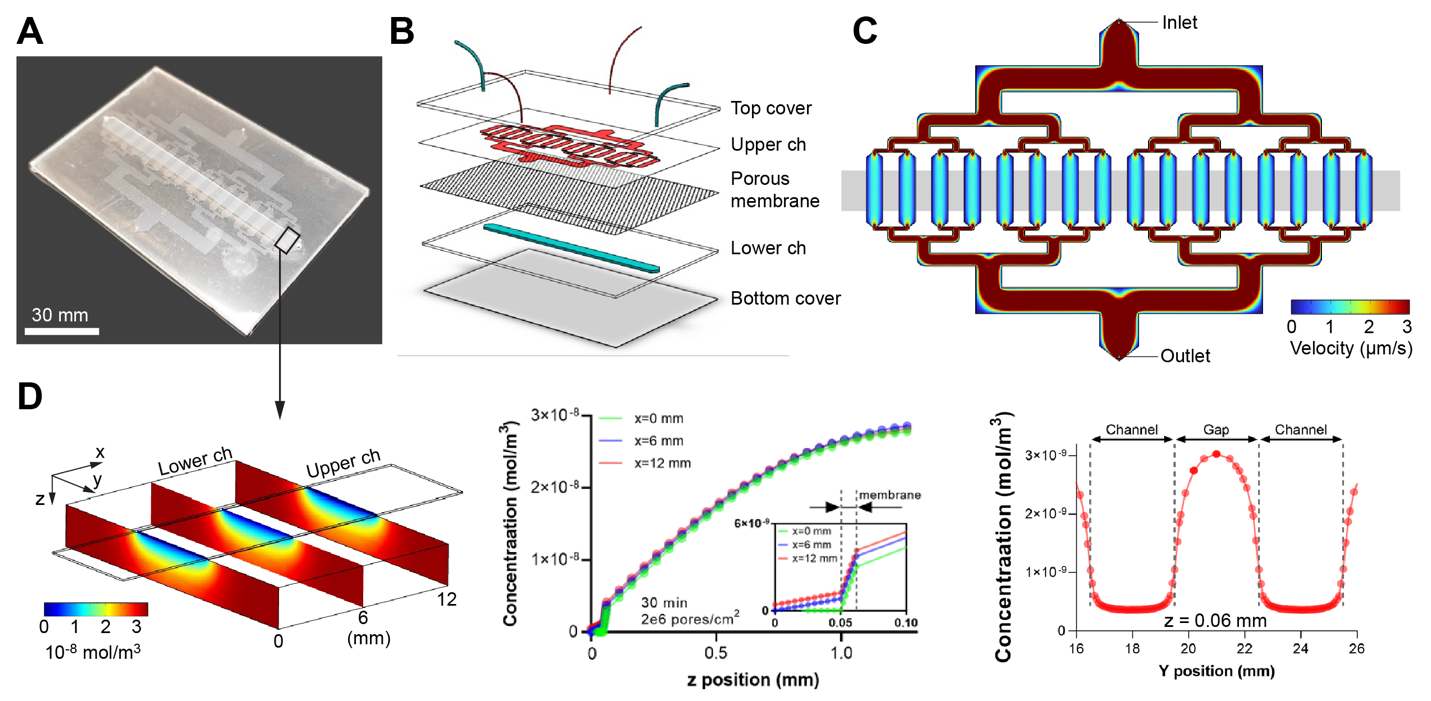

Fig. S1. T-Chip design enables stable and spatially uniform chemokine gradients across the porous membrane.

(A) Photograph of the fabricated transmigration-on-a-chip (T-Chip).

(B) Exploded schematic showing the T-Chip, consisting of an upper channel, porous membrane, and lower chemokine reservoir channel.

(C) Computational simulation of fluid velocity within the upper channel demonstrates uniform flow across the branched membrane regions.

(D) Computational simulation of CXCL10 diffusion in the lower channel beneath the membrane interface (boxed region in (A)) after 30 mins. 3D heatmap showing chemokine diffusion profile (left). Concentration gradient along the z-axis (height) showing consistent vertical gradients (middle). Chemokine concentration profile along the y-axis at z = 0.06 mm, confirming homogeneous lateral distribution (right).

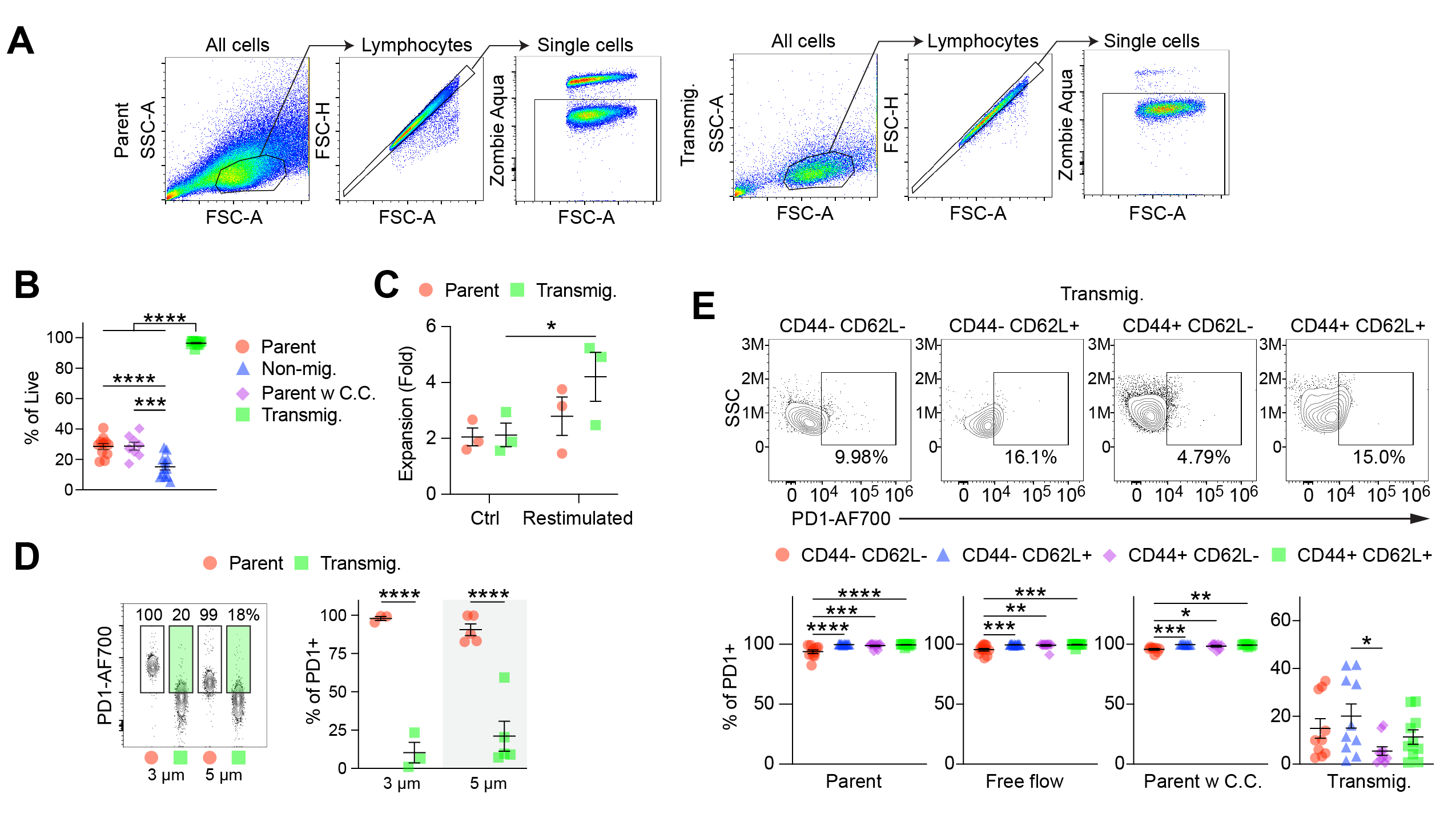

Fig. S2. Transmigration enriches for live CD8⁺ T cells and induces PD-1 downregulation independently of memory phenotype.

(A) Representative gating strategy for flow cytometry analysis of live CD8⁺ T cells from parent and transmigrated groups.

(B) Quantification of live cell percentages across parent, chemokine-exposed parent (Parent w. C.C.), non-migrated (perfused in T-Chip and collected from the upper channel), and transmigrated conditions. Transmigrated cells exhibited significantly higher viability compared to all other groups (n = 8-12 chips per group; mean ± SEM; ***p < 0.001, ****p < 0.0001, one-way ANOVA with Tukey’s multiple comparisons test).

(C) Expansion of parent and transmigrated CD8^+^ T cells measured 3 days after T-Chip processing under control conditions or following re-stimulation with Dynabeads (1:1 bead-to-cell ratio). Expansion fold was calculated based on total live cell numbers relative to the number of cells initially seeded for culture. (n = 3 chips per group; mean ± SEM; *p < 0.05, two-way ANOVA with Tukey’s multiple comparisons test).

**(D)** Representative flow cytometry plots (left) and summary (right) of surface PD-1 expression of CD8+ T cells transmigrated through 3 µm and 5 µm pores compared to parent cells, indicating acute loss of PD-1 irrespective of pore size. Green gates highlight the PD-1-low population observed following transmigration relative to the parent population. (n = 3–5 chips per group; mean ± SEM; ****p < 0.0001, two-way ANOVA with Tukey's multiple comparison test)

(E) PD-1 expression across memory subsets defined by CD44 and CD62L (n = 8-12 chips per group; mean ± SEM; *p < 0.05, **p < 0.01, ***p < 0.001, ****p < 0.0001, one-way ANOVA with Tukey’s multiple comparisons test). While parent and control groups maintained uniformly high PD-1 expression, transmigrated cells showed markedly reduced PD-1+ frequency across all subsets. This indicates that PD-1 downregulation is not restricted to a specific differentiation state. Representative PD-1 plots for each subset from the transmigrated group are shown on the right.

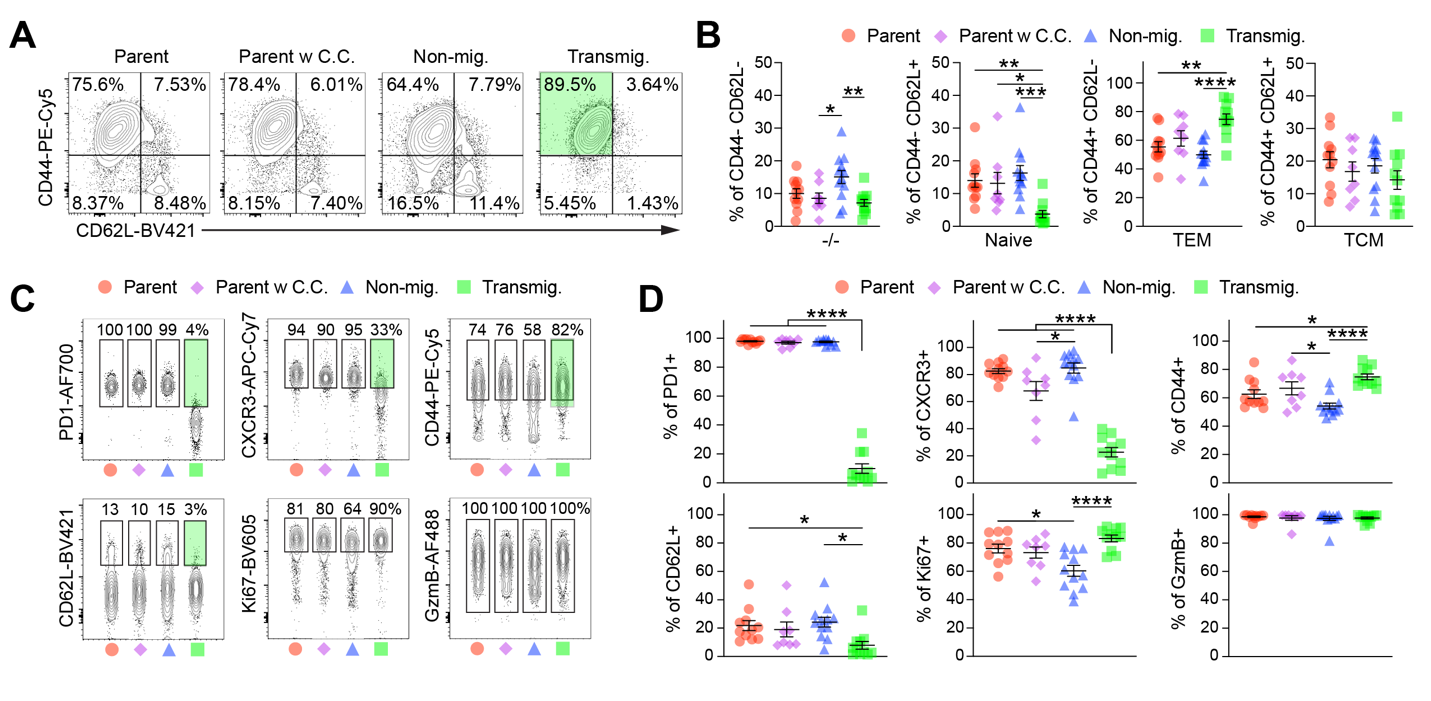

**Fig. S3. Transmigration through T-Chip reprograms CD8+ T cells by down-regulating PD-1 and enhancing effector phenotype and function.**

**(A)** Representative expression of PD-1, CXCR3, CD44, CD62L, Ki-67, and Granzyme B in parent CD8+ T cells, parent cells exposed to chemokine cues without flow (Parent w C.C.), non-migrated cells recovered from the upper channel (Non-mig.), and transmigrated cells recovered from the lower channel of T-Chip (Transmig.). Green gates highlight marker populations that were either enriched or reduced in transmigrated cells relative to the parent population. Full gating strategy for flow cytometry may be found in fig. S5A.

**(B)** Summary of marker-positive cells from (A). Transmigration significantly reduced PD-1+, CXCR3+, and CD62L+ populations while increasing CD44+ and Ki67+ populations. (n = 8-12 chips per group; mean ± SEM; *p < 0.05, **p < 0.01, ***p < 0.001, ****p < 0.0001, one-way ANOVA with Tukey's multiple comparison test)

**(C)** Representative CD44 vs. CD62L plots illustrating memory subsets in CD8⁺ T cells from each group. Green gates highlight the CD44⁺CD62L⁻ effector memory population enriched following transmigration.

**(D)** Summary of CD8+ T cell memory subsets defined by CD44 and CD62L expression: double-negative (-/-), naïve (CD44-CD62L+), effector memory (TEM; CD44+CD62L-), and central memory (TCM; CD44+CD62L+). Transmigration increased TEM cells and reduced naive subsets. (n = 8-12 chips per group; mean ± SEM; *p < 0.05, ****p < 0.0001, one-way ANOVA with Tukey's multiple comparison test)

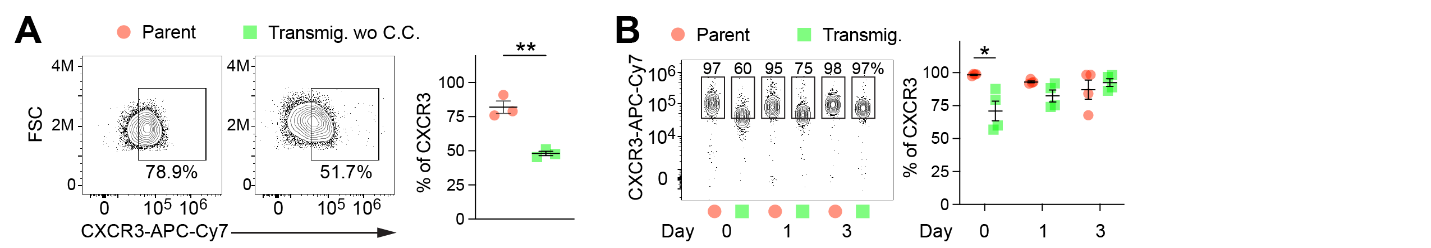

Fig. S4. Transmigration induces transient downregulation of surface CXCR3 on CD8⁺ T cells.

(A) Representative flow cytometry plots and quantification of surface CXCR3 expression in parent versus transmigrated CD8⁺ T cells processed in the absence of chemokine cues (Transmig. wo C.C.). Transmigration alone reduces surface CXCR3 levels, suggesting ligand-independent modulation (n = 3 chips per group; mean ± SEM; **p < 0.01, unpaired t-test).

(B) Time-course analysis of CXCR3 expression on parent and transmigrated cells. CXCR3 expression is gradually recovered to parental levels by 1-day post-transmigration (n = 4 chips per group; mean ± SEM; *p < 0.05, two-way ANOVA with Tukey’s multiple comparisons test).

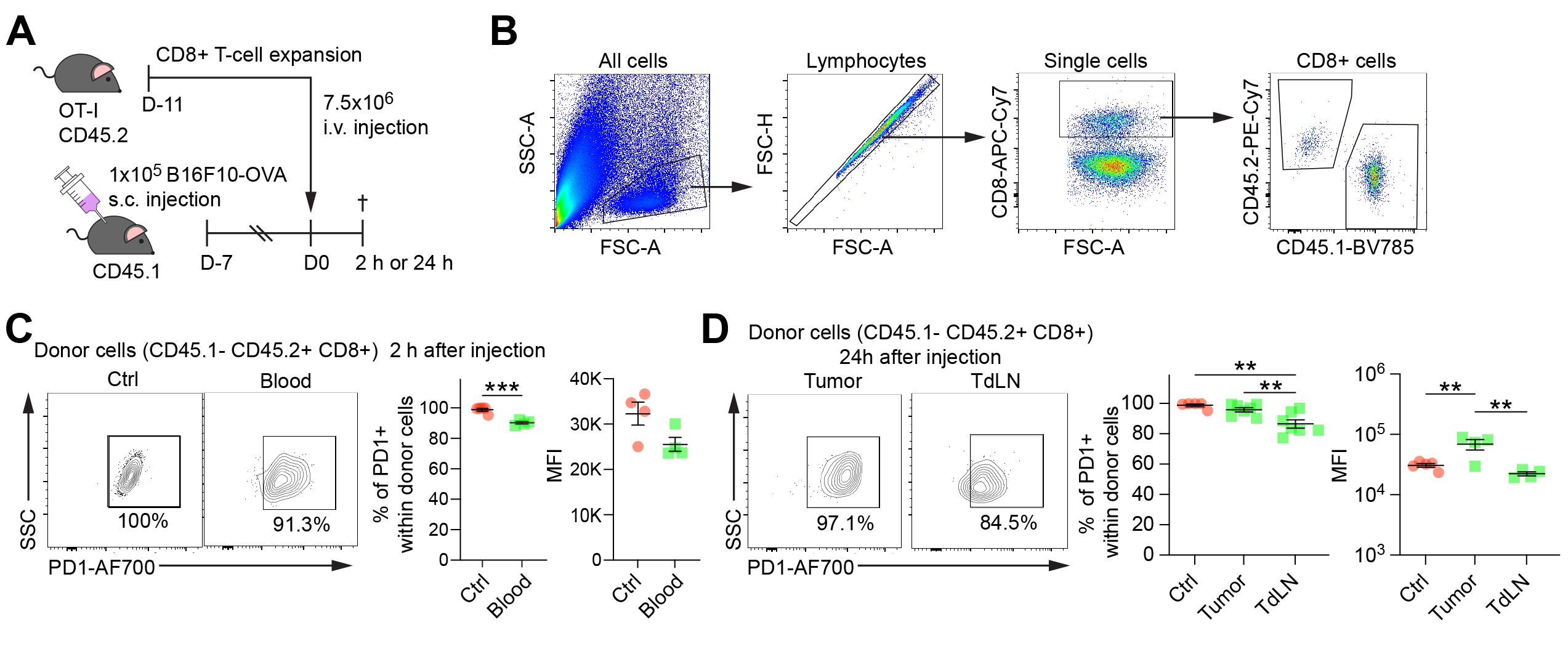

Fig. S5. PD-1 expression is largely maintained in circulation and recovers by 24 h after tissue homing.

(A) Experimental schematic for short-term adoptive transfer studies. Expanded OT-I CD45.2^+^ CD8^+^ T cells were injected intravenously (i.v.) into CD45.1^+^ mice bearing B16F10-OVA tumors, and donor cells were analyzed at 2 h or 24 h post-transfer.

(B) Gating strategy used for flow cytometry analysis of adoptive transfer experiment to assess early PD-1 regulation during tumor infiltration.

(C) Representative flow cytometry plots (left) and quantification (right) of PD-1 frequency and median fluorescence intensity (MFI) of donor CD8+ T cells (CD45.1^−^ CD45.2^+^) recovered from blood 2 h post-injection compared with non-injected control cells (Ctrl). (n = 4-5 mice per group; mean ± SEM; ***p < 0.001, unpaired t-test)

(D) Representative flow cytometry plots (left) and quantification (right) of PD-1 frequency and median fluorescence intensity (MFI) on donor CD8+ T cells (CD45.1^−^ CD45.2^+^) recovered from tumors and tumor-draining lymph nodes (TdLNs) 24 h after adoptive transfer. PD-1 expression was largely restored by 24 h in donor T cells recovered from tumors and TdLNs relative to the pre-transfer control (Ctrl) population. (n = 4-7 mice per group; mean ± SEM; **p < 0.01, one-way ANOVA with Tukey's multiple comparison test).

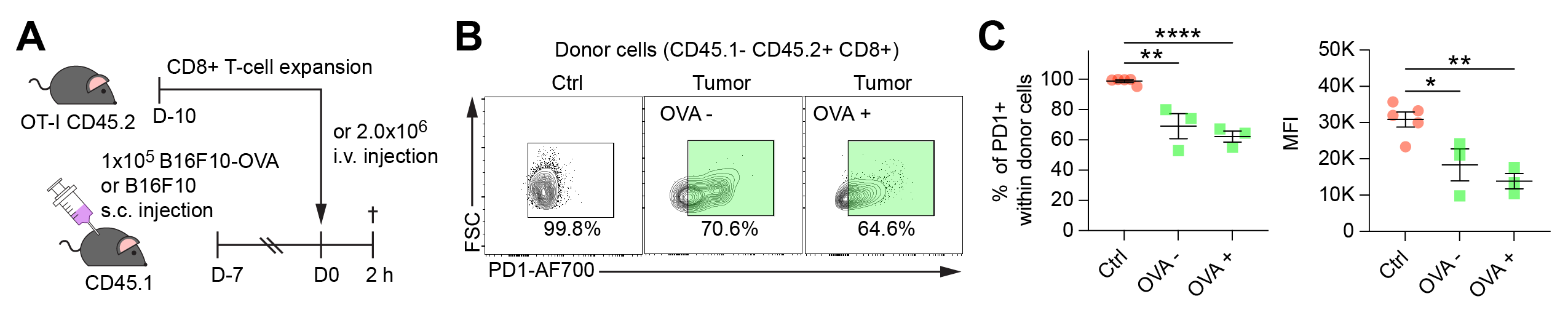

**Fig. S6. Acute PD-1 downregulation after intravenous transfer is independent of cognate antigen.**

**(A)** Experimental schematic for short-term adoptive transfer studies. Expanded OT-I CD45.2^+^ CD8^+^ T cells were injected intravenously into CD45.1^+^ mice bearing either B16F10-OVA or parental B16F10 tumors, and donor cells were analyzed at 2 h post-transfer.

**(B)** Representative flow cytometry plots and **(C)** quantification of PD-1 frequency and median fluorescence intensity (MFI) on donor CD8⁺ T cells (CD45.1⁻ CD45.2⁺) recovered from tumors of B16F10-OVA or parental B16F10 tumor-bearing mice 2 h after transfer. Green gates highlight the PD-1-low population observed following tumor homing relative to the pre-transfer control population. Acute PD-1 reduction at 2 h was observed in both tumor models, indicating that tumor OVA expression is not required for early PD-1 downregulation. (n = 3-5 mice per group; mean ± SEM; *p < 0.05, **p < 0.01, ****p < 0.0001, unpaired t-test).

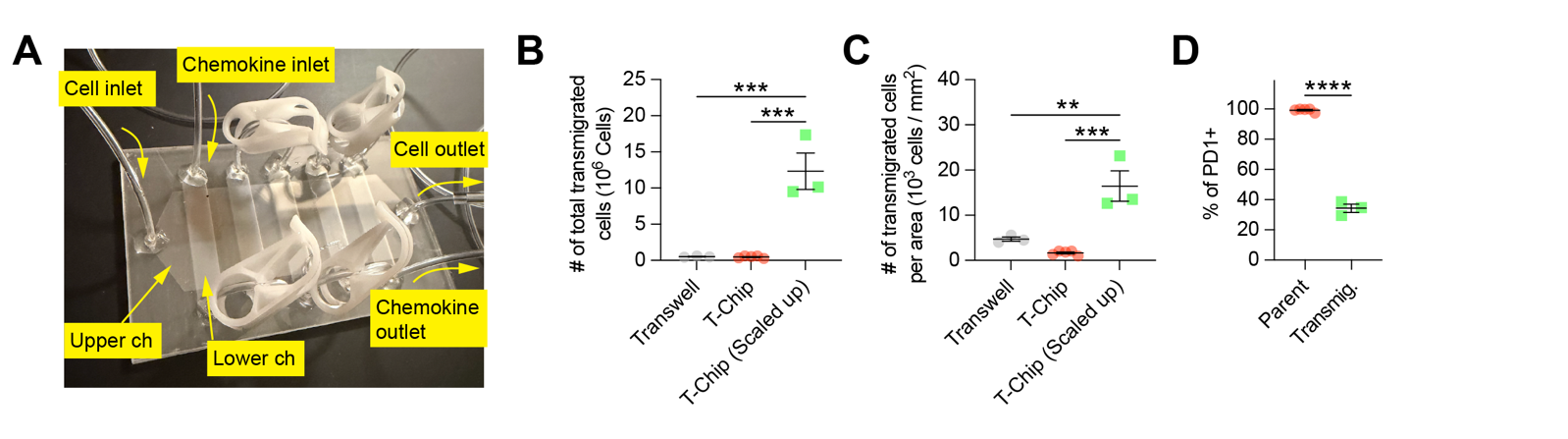

**Fig. S7. Scale-up T-Chip design substantially increases recovery of transmigrated CD8+ T cells while preserving transmigration-induced PD-1 loss.**

**(A)** Photograph of the scale-up T-Chip design with enlarged upper and lower channels and dedicated inlets and outlets for cell perfusion and chemokine loading.

**(B)** Quantification of total recovered transmigrated CD8+ T cells from conventional Transwell inserts, the original T-Chip, and the scale-up T-Chip design. (n = 3-5 wells or chips per group; mean ± SEM; ***p < 0.001, one-way ANOVA with Tukey’s multiple comparisons test).

**(C)** Number of recovered transmigrated CD8+ T cells normalized to the effective membrane area available for transmigration, demonstrating enhanced cell recovery efficiency of the scale-up T-Chip. (n = 3-5 wells or chips per group; mean ± SEM; **p < 0.01, ***p < 0.001, one-way ANOVA with Tukey’s multiple comparisons test).

**(D)** Surface PD-1 expression of parent and transmigrated OT-I CD8+ T cells processed in the scale-up T-Chip, showing that increased cell recovery is accompanied by preserved transmigration-induced PD-1 downregulation. (n = 2-5 mice per group; mean ± SEM; ****p < 0.0001, unpaired t-test).

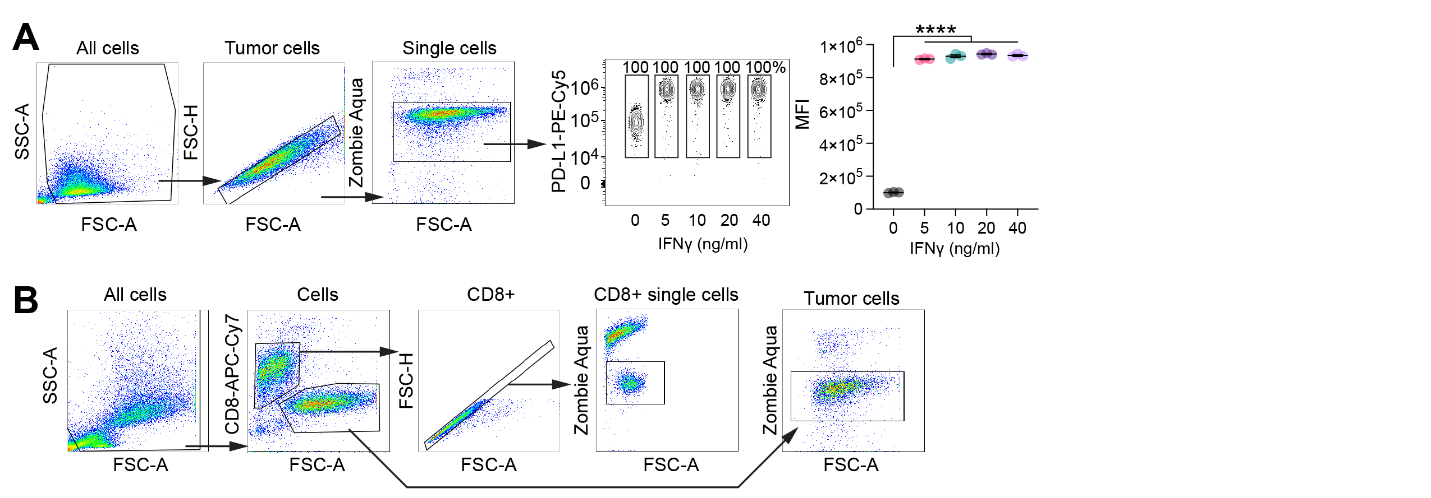

Fig. S8. IFN-γ induces PD-L1 upregulation in B16F10-OVA tumor cells used for co-culture assays.

(A) Gating strategy for flow cytometry analysis of live B16F10-OVA tumor cells and representative flow cytometry plots (left) and quantification of PD-L1 mean fluorescence intensity (MFI, right) on B16F10-OVA cells treated with increasing doses of IFN-γ (0–40 ng/mL). PD-L1 expression was significantly upregulated and reached saturation above 5 ng/mL (n = 3 per group; ****p < 0.0001, one-way ANOVA with Tukey’s multiple comparisons test).

(B) Gating strategy for analyzing CD8⁺ T cells and tumor cells recovered from co-culture, used for intracellular cytokine staining and tumor cell killing assays.

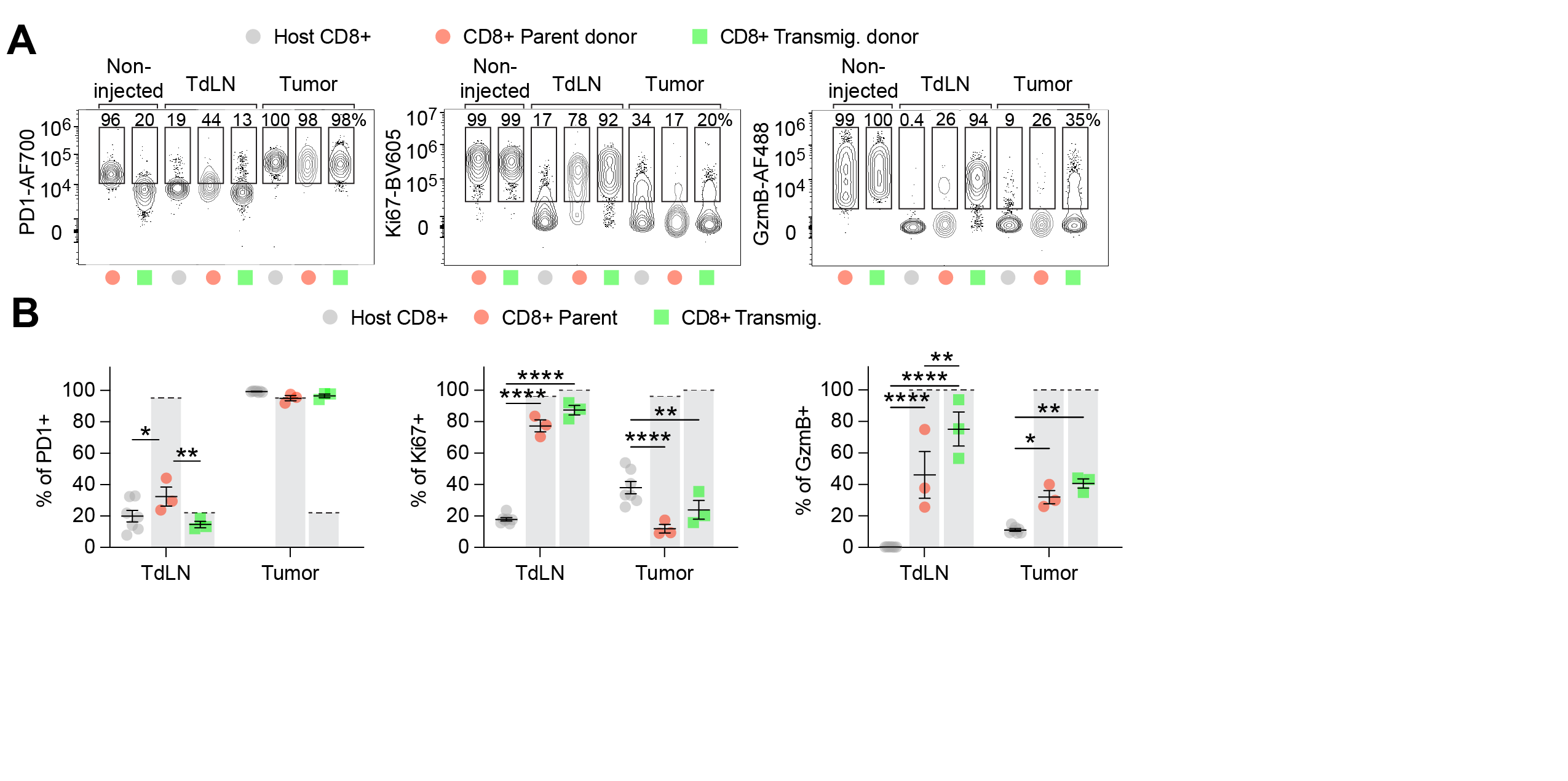

Fig. S9. Transmigrated donor CD8⁺ T cells maintain reduced PD-1 and elevated Ki-67 and GzmB in tumor-draining lymph nodes 7 days post-transfer. Following the experimental design in Fig. 3E–G, tumors were established on day 0, parent or T-Chip–transmigrated OT-I CD8⁺ T cells were administered peritumorally on day 7, and tissues were harvested 7 days later (day 14 after tumor implantation) for flow cytometry.

(A) Representative flow cytometry plots of PD-1, Ki-67, and GzmB expression in donor CD45.2⁺ CD8⁺ T cells recovered from tumors and TdLN.

(B) Quantification of PD-1⁺, Ki-67⁺, and GzmB⁺ frequencies in donor CD8⁺ T cells across tissues. (n = 3-7 mice per group; *p < 0.05, **p < 0.01, ****p < 0.0001, two-way ANOVA was used with Tukey's multiple comparison test).

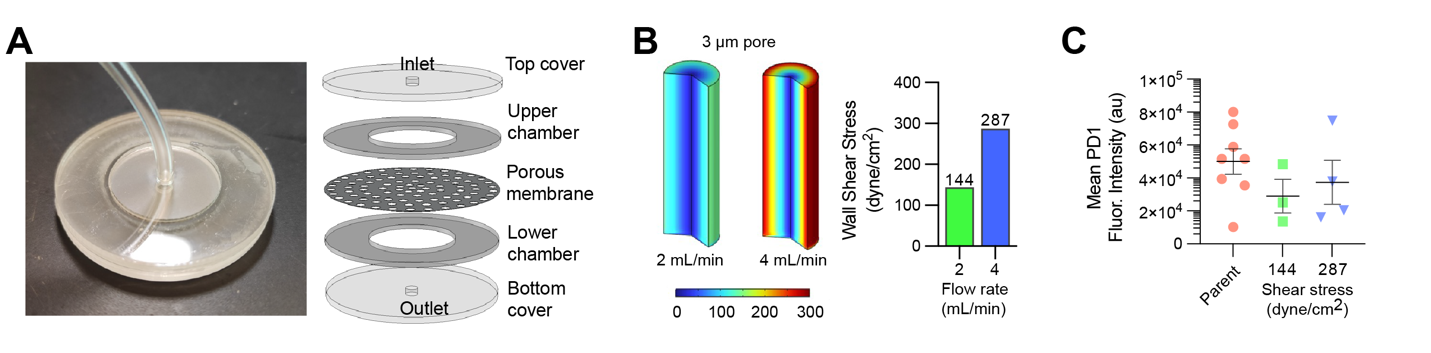

Fig. S10. Mechanical compression through pores does not induce PD-1 downregulation in CD8⁺ T cells.

(A) Photograph (left) and schematic (right) of a circular flow chamber used to apply high fluid pressure across a porous membrane. Cells were physically pushed against the membrane under high shear stress conditions.

(B) Computational modeling of wall shear stress at two input flow rates (2 and 4 mL/min), corresponding to 144 and 287 dyne/cm², respectively.

(C) Mean fluorescence intensity of surface PD-1 in CD8⁺ T cells subjected to mechanical compression across the membrane. No significant reduction in PD-1 was observed, indicating that passive mechanical pushing without transmigration is insufficient to induce PD-1 loss.

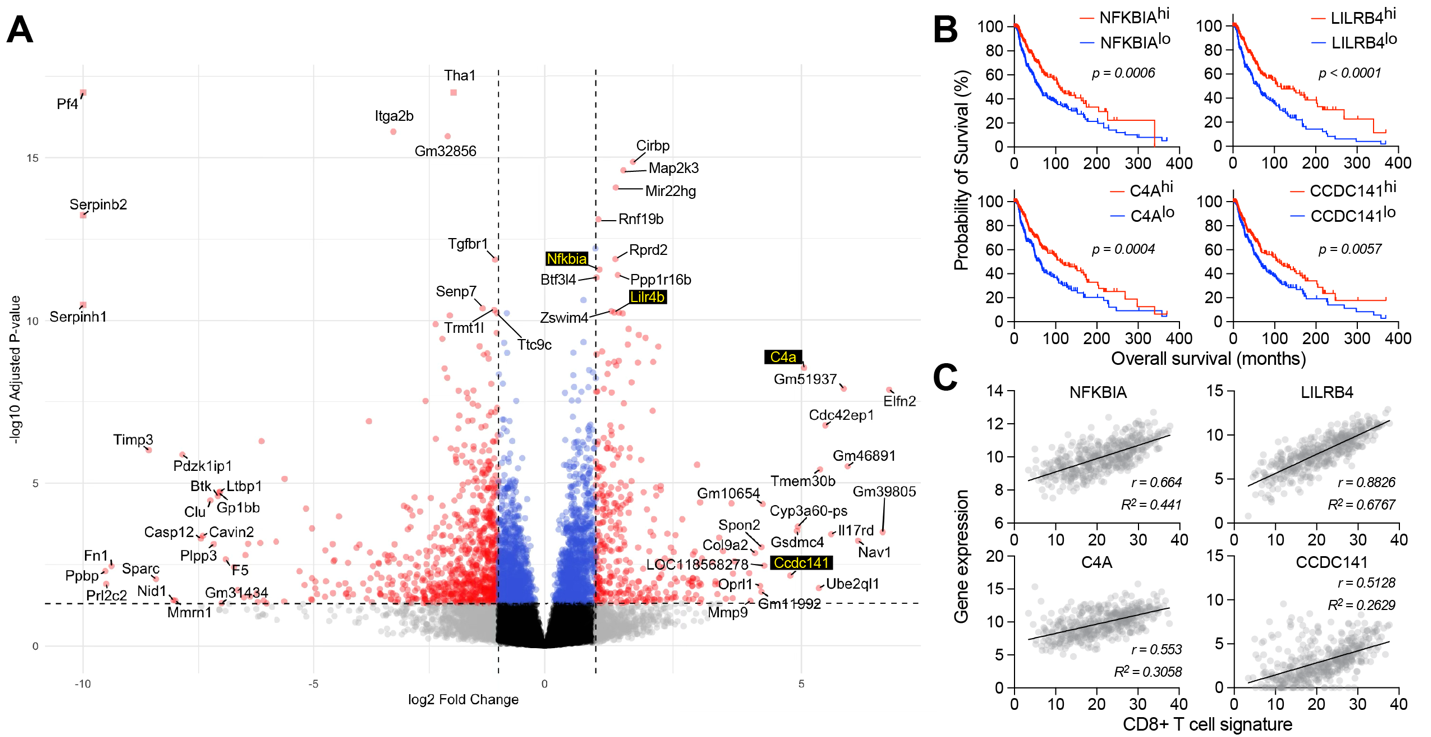

Fig. S11. Transmigration-associated DEGs include survival-linked genes in The Cancer Genome Atlas (TCGA) skin cutaneous melanoma (SKCM) cohort.

(A)  Extended volcano plot of bulk RNA-seq comparing parent and transmigrated mouse CD8+ T cells. The top 20 DEGs and the top 20 up- and down-regulated genes are labeled. Genes highlighted in yellow were associated with overall survival in the TCGA SKCM cohort.

(B) Kaplan–Meier survival curves for NFKBIA, LILRB4, C4A, and CCDC141 expression (high vs. low, median split) in the TCGA SKCM cohort, with log-rank P values indicated.

(C) Correlation of the same genes with a CD8+ T-cell signature score in the TCGA cohort. Gene expression values are shown as log₂(normalized count + 1). Lines indicate linear regression fits; r denotes correlation coefficient and R² the coefficient of determination.

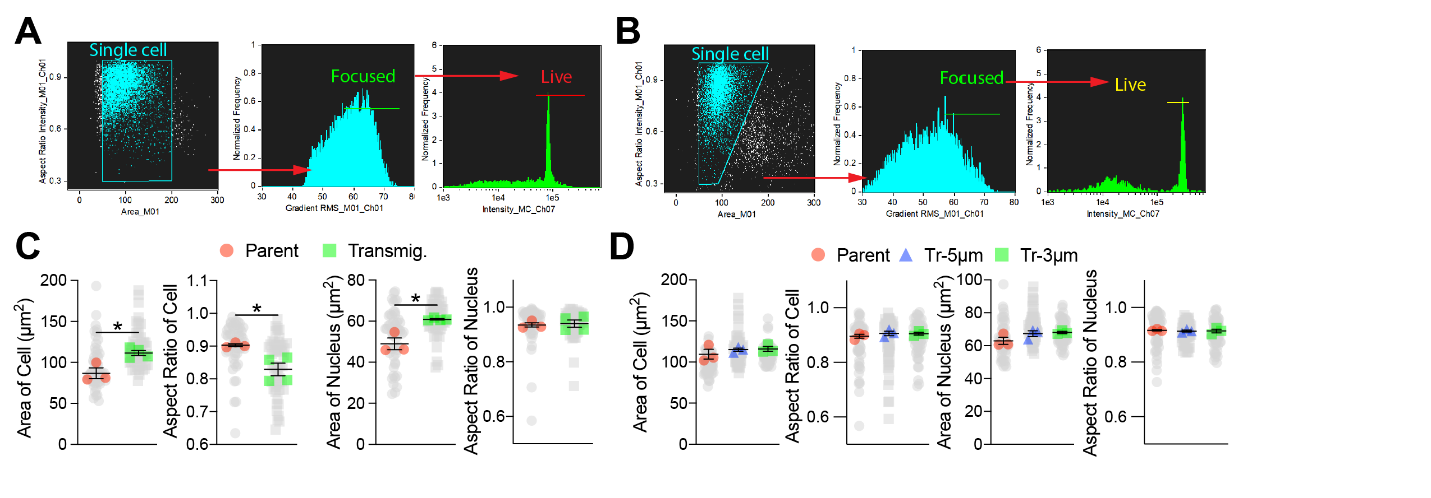

Fig. S12. Transmigration induces morphological remodeling in mouse but not human CD8⁺ T cells.

(A–B) Gating strategy for imaging flow cytometry analysis of mouse (A) and human (B) CD8+ T cells.

(C) Quantification of morphological parameters in mouse CD8+ T cells. Transmigrated cells exhibited increased cell area, reduced aspect ratio, and changes in nuclear size compared to parent cells (n > 1000 cells per group from 3 - 4 experiments; *p < 0.05, unpaired t-test). Gray dots indicate n = 50 randomly sampled individual cells to illustrate their distribution.

(D) Morphological analysis of human CD8⁺ T cells revealed no significant differences in cell body or nuclear geometry between parent and transmigrated groups (n > 1000 cells per group from 3 - 4 experiments). Gray dots indicate n = 50 randomly sampled individual cells to illustrate their distribution.

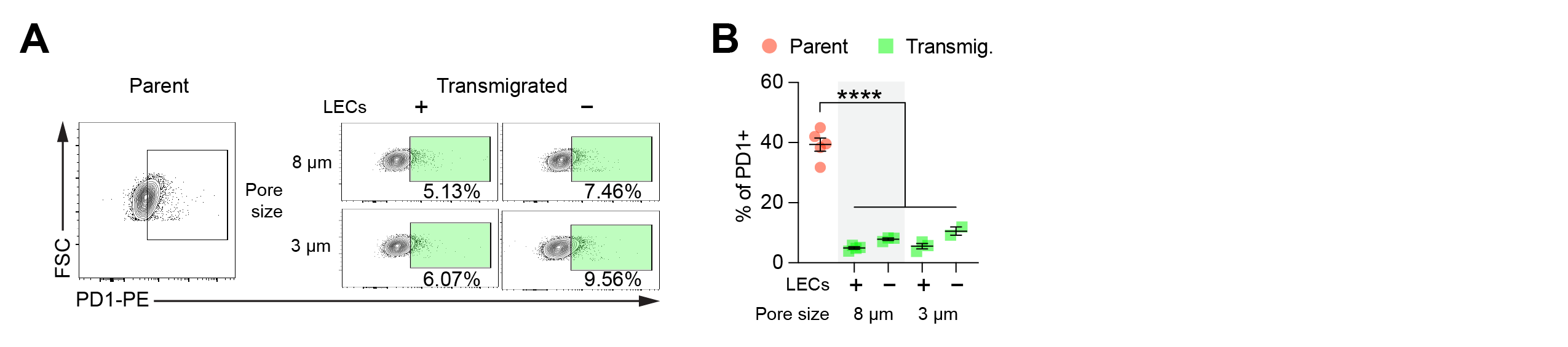

Fig. S13. Transwell assays with a lymphatic endothelial cell (LEC) monolayer reproduce transmigration-associated PD-1 downregulation in Human CD8⁺ T cells.

(A) Representative flow cytometry plots of PD-1 expression in parent and transmigrated human T cells recovered from Transwells seeded with a human LEC monolayer or membrane-only controls using 8 µm or 3 µm pore membranes. Green gates highlight the PD-1-low population observed in transmigrated cells.

(B) Quantification of PD-1⁺ frequencies across the indicated human Transwell conditions, showing PD-1 downregulation in transmigrated cells consistent with the murine assay. (n = 3-5 wells per group; mean ± SEM; ****p < 0.0001, one-way ANOVA with Tukey’s multiple comparisons test).

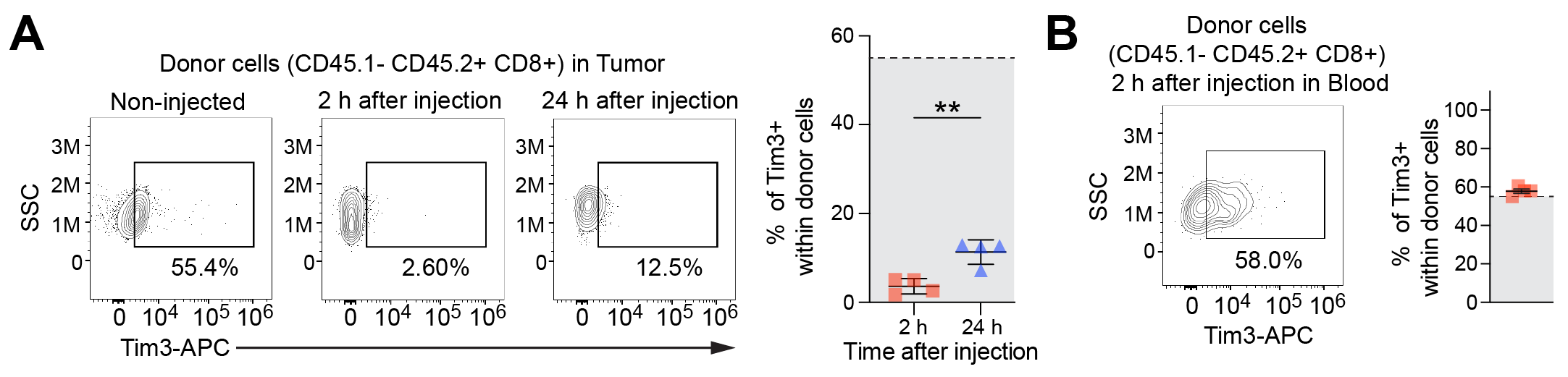

Fig. S14. Tim-3 is transiently downregulated during tumor infiltration but not observed from in vitro transmigration of expanded CD8 T cells.

(A) Tim-3 expression in donor OT-I CD8+ T cells (CD45.2+) recovered from tumors at 2 and 24 hours after adoptive transfer. CD8+ T cells were isolated from CD45.2+ OT-I mice, expanded ex vivo for 11 days, and intravenously injected into B16F10-OVA tumor-bearing CD45.1+ mice (1×10⁵ tumor cells implanted subcutaneously 7 days prior). Tumors were harvested at 2 h and 24 h post-injection for analysis. Tim-3 was markedly downregulated at 2 hours but partially restored by 24 hours, indicating a transient loss during early tumor infiltration (n = 4 mice per group; **p < 0.01, unpaired t-test).

(B) Tim-3 expression in donor CD8+ T cells recovered from blood at 2 hours post-transfer.

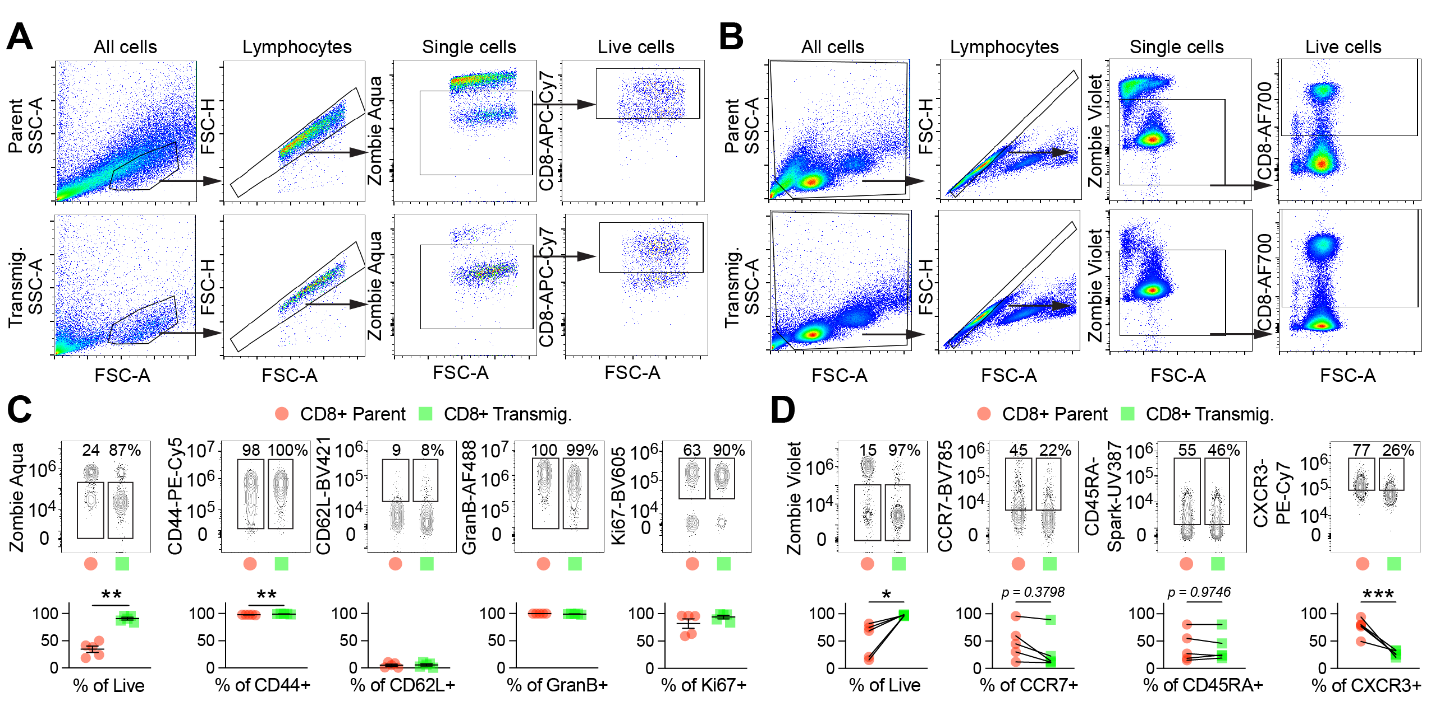

Fig. S15. Transmigration selectively enriches proliferative and viable subsets of CD8⁺ tumor-infiltrating lymphocytes (TILs) from both mouse and human tumors.

(A–B) Gating strategy for flow cytometry analysis of CD8⁺ TILs isolated from mouse (A) and human (B) tumors.

(C) In mouse CD8⁺ TILs, transmigration led to strong enrichment of live cells and increased frequency of CD44+ populations. Notably, even in samples with low baseline Ki-67⁺ frequency, transmigration enriched for highly proliferative Ki-67⁺ cells, suggesting selective retention of functionally active subsets. CD62L and Granzyme B levels remained unchanged (5 chips per group from 5 independent tumor fragments; **p < 0.01, unpaired t-test).

(D) Similarly, human CD8⁺ TILs isolated through transmigration showed robust enrichment of live cells from samples with variable viability, accompanied by selective downregulation of CXCR3, with CCR7 and CD45RA expression largely maintained (5 chips per group from 5 independent tumor fragments; *p < 0.05, ***p < 0.001, paired t-test).

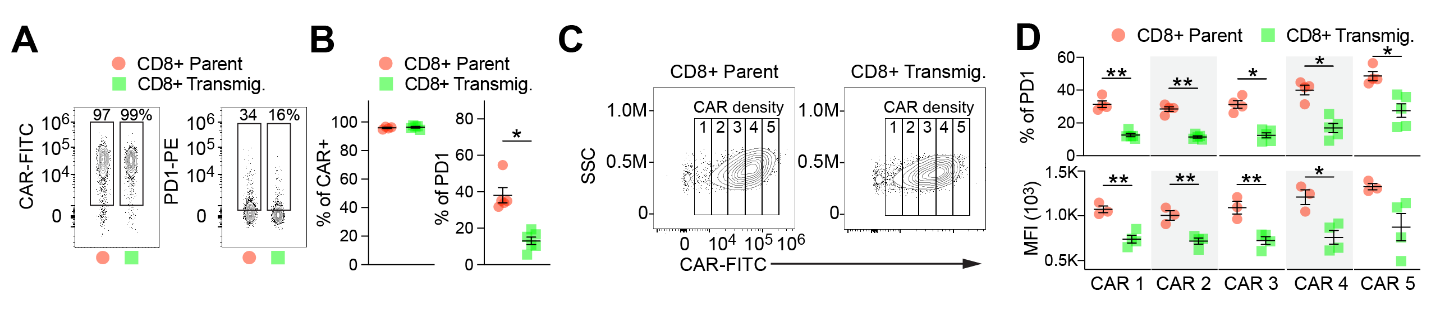

Fig. S16. Transmigration selectively reduces PD-1 expression in human CAR-T cells across a range of CAR expression levels.

(A) Representative flow cytometry plots of human CD8 CAR (anti-CD19) T cells showing CAR and PD-1 expression in parent and transmigrated populations.

(B) Quantification of CAR and PD-1 expression on CD8⁺ CAR-T cells. CAR expression was unaffected by transmigration, while PD-1+ frequency was significantly reduced (n = 4 - 6; *p < 0.05, unpaired t-test).

(C) Representative flow plots showing stratification of CAR signal into five density groups (CAR 1–5) in parent and transmigrated cells.

(D) PD-1 expression (top: % PD-1+; bottom: MFI) within each CAR density group. Transmigrated cells exhibited consistently lower PD-1 expression at all CAR levels, indicating that checkpoint regulation is uncoupled from CAR expression intensity (n = 4 per group; *p < 0.05, **p < 0.01, unpaired t-test).

| **Antibodies** | **Source** | **Identifier** |
| --- | --- | --- |
| BV785 anti-mouse CD45.1 Antibody | Biolegend | Cat#110743 |
| PE/Cyanine7 anti-mouse CD45.2 Antibody | Biolegend | Cat#109830 |
| APC/Cyanine7 anti-mouse CD45.2 Antibody | Biolegend | Cat#109823 |
| APC/Cyanine7 anti-mouse CD8a Antibody | Biolegend | Cat#100714 |
| Spark UV 387 anti-mouse CD8a Antibody | Biolegend | Cat#100797 |
| Alexa Fluor 700 anti-Mouse PD-1 Antibody | R&D Systems | Cat#FAB7738N |
| APC anti-mouse CD366 (Tim-3) Antibody | Biolegend | Cat#119706 |
| APC/Cyanine7 anti-mouse CD183 (CXCR3) Antibody | Biolegend | Cat#126552 |
| PE/Cyanine5 anti-mouse/human CD44 Antibody | Biolegend | Cat#103010 |
| BV421 anti-mouse CD62L Antibody | Biolegend | Cat#104436 |
| BV605 anti-mouse Ki-67 Antibody | Biolegend | Cat#652413 |
| Alexa Fluor 488 anti-human/mouse Granzyme B Recombinant Antibody | Biolegend | Cat#396424 |
| BV785 anti-mouse IFN-γ Antibody | Biolegend | Cat#505837 |
| BV605 anti-mouse IL-2 Antibody | Biolegend | Cat#503829 |
| Alexa Fluor 488 anti-mouse TNF-α Antibody | Biolegend | Cat#506313 |
| PE/Cyanine5 anti-mouse CD274 (B7-H1, PD-L1) Antibody | Biolegend | Cat#124344 |
| PE Mouse Anti-Human CD279 (PD-1) Antibody | BD Biosciences | Cat#557946 |
| PE/Cyanine7 anti-human CD183 (CXCR3) Antibody | Biolegend | Cat#353719 |
| Spark UV 387 anti-human CD45RA Antibody | Biolegend | Cat#304179 |
| BV 785 anti-human CD197 (CCR7) Antibody | Biolegend | Cat#353230 |
| BV605 anti-mouse CD279 (PD-1) Antibody | Biolegend | Cat#135219 |
| *InVivo*MAb anti-mouse CD16/CD32 | BioXcell | Cat#BE0307 |
| FITC-conjugated anti-idiotype antibodies, clone 19E3 (CD19-targeted), | Memorial Sloan Kettering Cancer Center Antibody and Bioresource Core Facility | - |
| CellTrace™ Blue Cell Proliferation Kit, for flow cytometry | Invitrogen | Cat#C34568 |
| Zombie Violet™ Fixable Viability Kit | Biolegend | Cat#423113 |
| Zombie Aqua™ Fixable Viability Kit | Biolegend | Cat#423102 |

Table S1. List of antibodies used for flow cytometry analysis.

Movie S1.

Real-time bright field imaging of mouse CD8⁺ T cells migrating through 3 µm-pore in T-chip.

Movie S2 & S3.

Real-time confocal imaging of pre-stained PD-1 (Red) loss in CD8⁺ T cells migrating through a horizontal confined microchannel (width × height = 5 µm × 5 µm), with nuclei labeled in blue.
